# Large-scale Perturbation of Systems Biology-Derived Genes Reveals Modifiers of HD-associated Transcriptomic Networks and Pathology

**DOI:** 10.64898/2026.03.02.709091

**Authors:** Peter Langfelder, Nan Wang, Lalini Ramanathan, Young Mi Oh, Seongwon Lee, Fuying Gao, Xiaofeng Gu, Matthew Stricos, Mary Plascencia, Raymond Vaca, Jeffrey Richman, Thomas F. Vogt, Steve Horvath, Andrew S. Yoo, Jeffrey Aaronson, Jim Rosinski, X. William Yang

## Abstract

In Huntington’s disease (HD), disease-driver genes are broadly expressed, but pathogenic players specific to vulnerable neurons remain poorly defined. Leveraging a previously defined mutant huntingtin (mHtt) CAG-length-associated gene network, we perturbed 115 module hub genes with heterozygous knockout (KO-het) to assess their striatal transcriptomic modifier effects in wildtype and Q140 HD mice. We generated 3,592 striatal RNA-seq datasets, mapped 6,517 perturbagen-responder gene pairs, and uncovered regulators of medium spiny neuron (MSN) identity gene expression and DNA-methylase/demethylase-sensitive genes. We developed a bioinformatic pipeline to rank the perturbations with significant impacts on striatal transcriptome and HD-associated gene networks in wildtype or Q140 mice. KO-het for FoxP1 and Scn4b (two MSN-selective genes) exacerbated, whereas Pdp1 KO-het ameliorated, Q140 striatal pathology. Importantly, knockdown of functionally opposing ion channels, SCN4B and KCNH4, dichotomously affected aggregation and neurodegeneration in reprogrammed HD patient MSNs. Together, our study rigorously evaluated systems biology-derived candidates to identify modifiers of HD-associated molecular networks and pathology, providing an in vivo perturbation- transcriptome resource and highlighting genes involved in MSN excitability, transcription, calcium signaling, and mitochondrial metabolism in HD pathogenesis.

## INTRODUCTION

Huntington’s disease (HD) is a dominantly inherited neurodegenerative disorders caused by an expanded CAG repeat (≥ 40 repeats) in Huntingtin (HTT) gene. The onset of HD, defined by its motor symptoms, is usually around middle age but can span from childhood to advanced age. HD age at onset is inversely correlated with the length of the inherited mutant CAG alleles (Lee et al., 2019). HD is clinically characterized by a triad of motor, cognitive, and psychiatric symptoms, and its disease course is a progressive decline with premature death about 15-20 years after onset (Vonsattel & DiFiglia, 1998). HD neuropathology is characterized by massive degeneration of striatal medium spiny neurons (MSNs), including both D1- and D2-MSN types. There are also cortical atrophy and degeneration of deep-layer cortical pyramidal neurons (Fu et al., 2018). Additionally, a pathognomonic neuropathological marker of HD is aggregation of mutant Huntingtin (mHTT) protein, which can be in the nucleus as well as the neuropil (Gutekunst et al., 1999). Currently, there are no disease-modifying therapies available for HD.

How mHTT elicits selective neuronal pathogenesis, particularly in vulnerable MSNs, remains an enigma. Mutant HTT is broadly expressed in the brain and peripheral tissues at all ages, and it has hundreds if not more potential protein interaction partners involved in a wide range of functions including but not limited to vesicular trafficking, cytoskeleton organization, synaptic transmission, RNA binding and processing, and chromatin and transcriptional regulation (Culver et al., 2012; Greco et al., 2022; Shirasaki et al., 2012; Stroedicke et al., 2015). To date, there is no proven striatal MSN-selective function of HTT that can be linked to HD neuronal vulnerability. Another mechanism, somatic expansion of the mutant CAG repeats, has been strongly implicated in selective neuronal pathogenesis in HD. In HD postmortem brains, the vulnerable striatal MSNs (Pressl et al., 2024; Handsaker et al., 2025) and deep layer cortical neurons (Pressl et al., 2024), but not the vast majority of non-vulnerable cell types, show more prominent somatic CAG repeat expansion and transcriptionopathy (Matlik et al., 2024; Pressl et al., 2024). Moreover, Genome-wide Association Studies (GWAS) of modifiers of HD age-at-onset identify significant impacts of mismatch repair (MMR; i.e. MSH3, PMS1 and MLH1) and other DNA repair genes (Genetic Modifiers of Huntington’s Disease, 2015; Genetic Modifiers of Huntington’s Disease Consortium. Electronic address & Genetic Modifiers of Huntington’s Disease, 2019). Importantly, knockout of distinct MMR genes, particularly Msh3 and Pms1, greatly reduces the MSN-selective somatic CAG repeat expansion, transcriptomic dysregulation, and mHtt aggregation in Q140 knockin mice (Wang et al., 2025). These studies convergently show that not only mHTT but also distinct MMR complex genes drive selective neuronal pathogenesis in HD. However, since MMR genes and proteins are nearly ubiquitously expressed, similar to mHTT, it remains unclear how MSN-specific biological processes intersect with these known disease drivers to elicit selective neuronal pathogenesis in HD.

Systems biology has become a mainstay in organizing and interpreting complex omics datasets (Parikshak et al., 2015). One powerful systems biology tool to analyze large gene expression datasets is Weighted Gene Co-expression Network Analysis (WGCNA; Langfelder and Horvath, 2008), which can identify groups of genes (modules) with highly correlated expression patterns across datasets. The advantage of WGCNA analysis is that it reduces the high dimensionality of the entire transcriptomic dataset down to dozens of coherently expressed gene modules, with each module having “hub genes” that are highly connected (i.e. correlated) with other genes in the module and hence are the most representative of the distinct expression profiles of a given module. Although these WGCNA module hub genes are often used in gene set enrichment analysis to identify enriched biological pathways or other molecular features (e.g. diurnally expressed genes or disease susceptibility genes; Wang et al., 2022; Hartl et al., 2021), to our knowledge, there are no studies the mammalian system that have systematically performed in vivo genetic perturbation of hub genes for their impacts on gene expression and other phenotypes.

Our prior study of large transcriptomic datasets from an allelic series (AS) of HD mouse models - carrying six inherited mHtt CAG repeat lengths ranging from 20 to 175 - defined 13 WGCNA modules strongly negatively (5 modules) or positively (8 modules) associated with inherited mHtt CAG lengths (Langfelder et al., 2016). Among these modules, the top CAG- downregulated module (M2 with 1,867 genes) is enriched for genes selectively expressed in MSNs (i.e. MSN identity genes) and other neuronal and synaptic genes, while the top CAG-upregulated module (M20; 838 genes) is enriched with genes involved in cell division and DNA damage response. Additionally, we also performed striatal proteomic studies of the AS mice, which helped define mHtt polyglutamine length (“Q-length) dependent proteins and WGNCA protein modules (Langfelder et al., 2016). Moreover, our recent genetic study of HD GWAS/MMR genes showed that the mHtt CAG-dysregulated genes and modules are highly selective for striatal MSNs, depend on MMR-driven somatic repeat expansion, and significantly overlap with alterations observed in HD patient MSNs (Wang et al., 2025; Gu et al., 2022; Handsaker et al., 2025; Matlik *et al*, 2024). However, it remains unclear whether mHtt-dysregulated genes may have functional roles in striatal neuronal pathogenesis in HD.

In this study, we hypothesized that the hub genes from the AS Htt CAG- and Q-length dependent modules (Langfelder *et al*., 2016) are a rich source of in vivo modifiers -including MSN-selective regulators - of HD-associated transcriptome and pathology. To rigorously test this idea, we selected 115 hub genes from these modules and built a streamlined, generalizable, and scalable pipeline to analyze and rank genetic modifier effects on the HD-associated striatal transcriptome and gene networks. Our study identifies both MSN-selective and broadly expressed genes that exert protective or pathogenic effects on HD-associated gene expression and neuropathology, highlighting genes involved in MSN excitability, transcription, Ca^2+^ signaling, and mitochondrial metabolism.

## RESULTS

### A scalable germline heterozygosity KO perturbation pipeline to test modifiers of HD transcriptomic networks in vivo

To select top HD gene signature related genes for in vivo perturbations, we used a combination of statistics that include: (i). DE gene Z statistics for gene association with mHtt CAG length’ (ii). AS CAG-dependent module membership Z statistics; (iii). and module eigengene association Z statistics with CAG length to produce ranking of top genes in the analysis (See Methods for details). This initial criterion is dominated by the hub genes in AS M2 module (68 genes) but also include genes in M20 (7 genes), M1 (13 genes), and M7 (2 genes). Additionally, we selected 10 genes from a CHDI/GNS causal modeling of AS transcriptomes (www.hdinhd.org), pM2 protein networks (3 genes), and other striatal specific transcription factors or chromatin factors (19 genes). Importantly, among our selected gene list there are multiple genes previously found at GWAS modifier loci (Gsg1l, Srebf2), genes (Hpca, Pde10a) involved in other movement disorders such as dystonia (Charlesworth et al., 2015), or numerous genes previously implicated in HD molecular pathogenesis (e.g. Rest, Itpr1, Pde10a, Rasd2/Rhes, Tcf20, Gpx6) (Zuccato et al., 2003; Tang et al., 2003; Beaumont et al., 2016; Subramaniam et al., 2009; Yamanaka et al., 2014; Shema et al., 2015). However, the disease-modifying roles of these genes have not been evaluated in a full-length mHtt KI mouse model in vivo.

We decided to use germline heterozygous KO alleles to test their effects in modifying striatal transcriptome in WT or HD mouse backgrounds (Figure 1A). This is to avoid the caveats of using homozygous mutants, such as potential lethality or developmental phenotypes and high cost and poor scalability. We crossed KO-het alleles of 115 genes (Supplemental Table S1) with heterozygous Q140 mice carrying an expanded ∼140 CAG trinucleotide repeats in the endogenous murine Htt gene (Menalled et al., 2003; Wang et al., 2025). We obtained 82 KO alleles from existing mutant mouse repositories (see Methods), and importantly, also employed CRISPR/Cas9 to create 33 novel KO-het alleles in this and a prior study (Wang et al., 2022). For each KO gene allele, we generated four genotypes (WT, Q140, KO-het, and Q140/KO-het; N=8 per genotype, sex balanced) and aged them to 6-months (denoted as 6m) before dissecting the striatal tissues for bulk RNA-seq (Wang et al., 2022). We obtained a total of 3,592 striatal RNA-seq samples, an aggregate of 903 WT samples, 900 Q140 samples, and 896 KO-het and 893 Q140/KO-het samples of different genotypes. For each RNA-seq sample, we detect between 16,557 and 17,372 genes (with an average of 17,149 genes). Together, our study represents - to our knowledge - one of the germline KO - transcriptomic perturbation datasets in the mammalian brain, providing a rich resource to discern the underlying striatal biology and HD pathogenesis.

**Figure 1.**
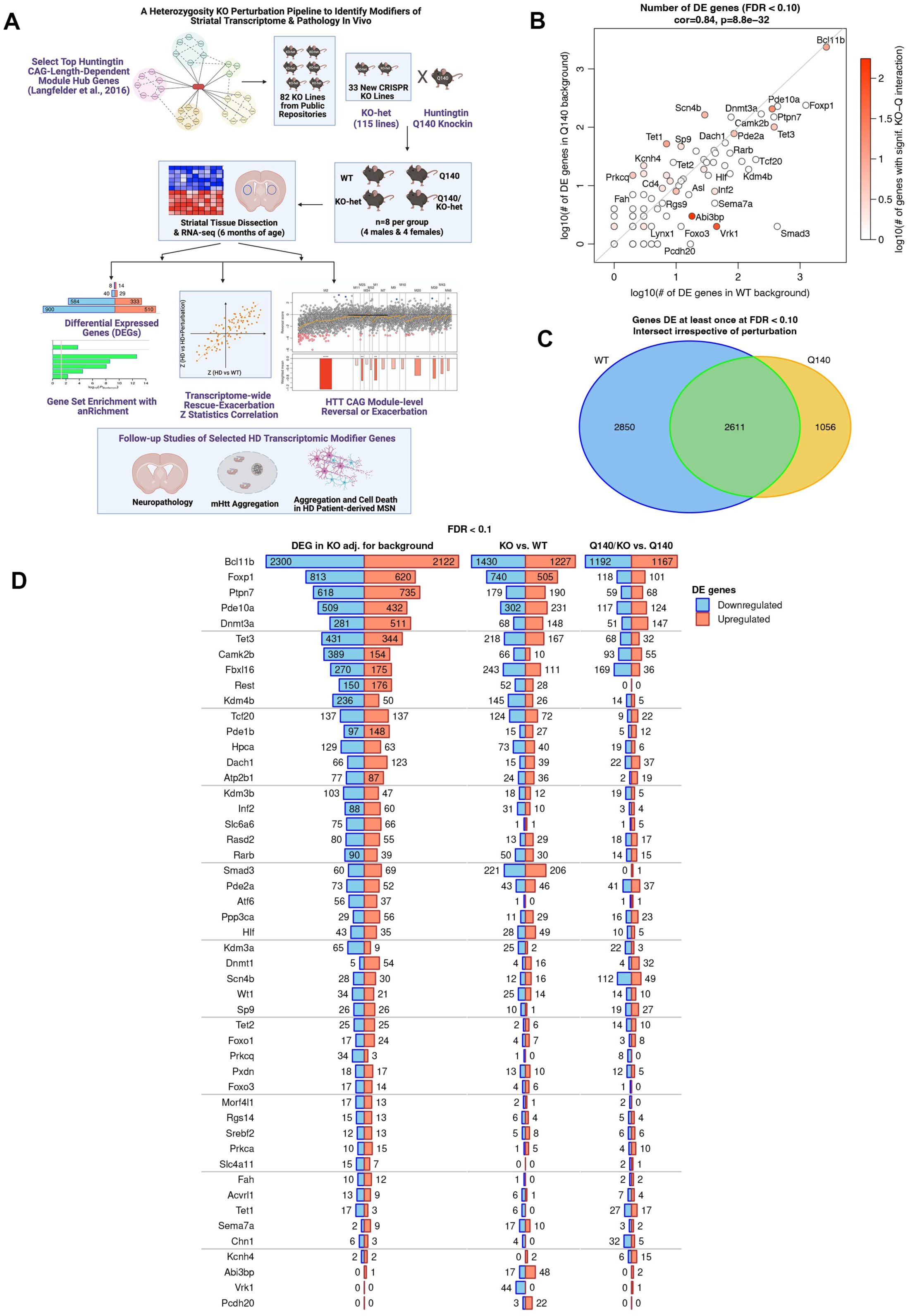
Germline heterozygous knockout perturbation and striatal transcriptomic analyses of 115 genes in WT and Q140 HD knockin mice. (A) Schematic diagram of the main elements of our study. (B) Scatterplot of log-transformed numbers of DE genes in WT (x-axis) and Q140 (y-axis) backgrounds. Each point represents one KO-het. Color represents the number of genes with significant Q140-KO-het interaction. (C) Venn diagram representing numbers of genes DE at FDR<0.1 in WT (blue) and Q140 (orange), and the number of genes DE in the same direction for the same KO-het in both WT and Q140 backgrounds. (D) Numbers of genes DE at FDR<0.1 in analysis across both backgrounds with background as a covariate, in WT background only and in Q140 background only. KO-het perturbations with at least 20 DE genes in one of the analyses are shown.

### Overview of the differential gene expression analysis of KO-het perturbations in WT and Q140 background

We first carried out differential expression (DE) analysis among the 4 genotypes for each cross and identify KO-het associated differential expressed genes (DEGs; FDR < 0.1) in either WT or Q140 backgrounds (Figure 1C-1D). Among these, there are 50 KO-het perturbations that elicit at least 20 DEGs in either WT, Q140, or KO adjusted for background (Figure 1D; Supplemental Table S2; see Methods). We found numbers of DEGs in WT and Q140 backgrounds are strongly correlated across the perturbations (r = 0.82, p=6.3e-28, Figure 1B). However, the number of DEGs in WT background tends to be higher than those in Q140 background for many KO-hets (Figure 1B, 1D). Among those with the largest number of DEGs, they are either transcription or chromatin factors (Bcl11b, Foxp1, Tcf20, Dnmt3a, Tet3, Kdm3b) or signaling proteins (Ptpn7, Pde10a, Pde2a, Camk2b). Together, we have established a dataset of KO-het perturbations and resulting DEGs (FDR<0.1) that contain 6517 genes, with 2611 shared between WT and Q140 backgrounds in the same perturbations (Figure 1C; Supplemental Table S3). In a DE analysis across adjusted for background we identify 8357 genes with significant KO-het effects (Supplemental Table S3). This ensemble of perturbagen-responder gene set can be readily searched to further explore perturbed genes with shared upstream regulators (Supplemental Figure S1), which may facilitate the study of specific genes of interest in the striatal biology and disease.

We next performed enrichment analysis of the ensemble gene set using an in-house gene enrichment R package that incorporates publicly available gene sets (e.g. GO, KEGG, Reactome, NCBI Biosystems, MSigDB, ChEA), HD-related gene sets (www.HDinHD), as well as our internally curated gene sets for brain cell types, striatal biology, and WGCNA modules see Methods). This analysis in the WT or Q140 perturbation gene sets reveal significant enrichment terms (Supplemental Table S4; Supplemental Figure S2), and the top ones in WT background include the following categories: genes downregulated or upregulated in striatal allelic series mice (Q140, Q175); Top D1-MSN or D2-MSN marker genes; common MSN marker genes; Genes down- or up-regulated in Foxp1 KO mice; Synaptic genes (GO); and Signaling by GPCR (Reactome).

We clustered top enriched terms in top 20 KO-hets in either WT (Figure 2A) or Q140 (Figure 2B) backgrounds based on similarity of their enrichment across the KO-hets (Methods). This analysis in WT background reveals a set of genes regulate the expression of genes enriched in specific striatal neuronal and glial cell types (Figure 2A). Importantly, a cluster of KO-hets either downregulate (Hpca, Bcl11b, Foxp1, Rarb, Camk2b, and Fbxl16) or upregulate (Vrk1, Abi3bp, Dach1, Pde10a) D2-MSN enriched genes. Several KO genes also downregulate (Rarb) or upregulate (Hfl, Bcl11b) D1-MSN genes. The roles of Foxp1 and Rarb in regulating MSN-type-specific gene expressions are shown before (Anderson et al., 2020; Rataj-Baniowska et al., 2015), validating the robustness of our datasets. Interestingly, a prior striatal FoxP1 DEG sets are also significantly overlap with those of Vrk1, Pde10a, Ptpn7, Tcf20, Pde2a, Hpca, Bcl11b, Camk2b and Fbxl16, suggesting potential shared - and likely MSN-related - gene regulation. In Q140 backgrounds, top enriched terms remain to be striatal cell-type specific genes, in addition to FoxP1 KO as well as various HD mouse gene signatures (Figure 2B). Interestingly, several genes appear to selectively downregulate MSN-specific genes in Q140 but not WT background (i.e. Scn4b and Sp9), with Sp9 being specific to D2-MSN genes. Another surprising finding is that Fbxl16 appears to selectively downregulate oligodendrocyte genes in Q140 background but MSN genes in WT background.

**Figure 2.**
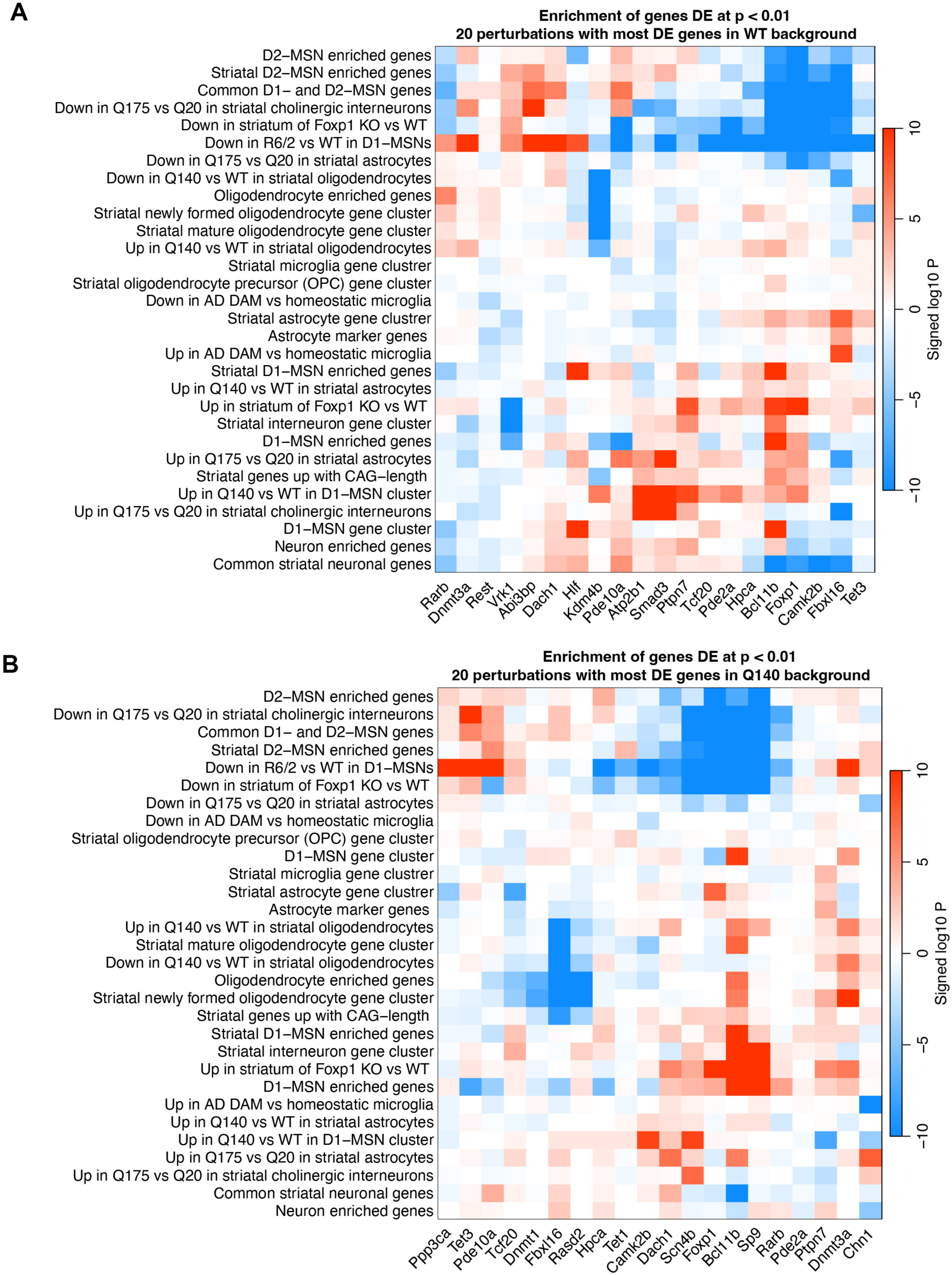
Clustering of top enrichment terms for the top 20 heterozygous knockout perturbations in WT and Q140 mice. (A) Enrichment heatmap of genes DE at p<0.01 for KO-het vs. WT in the 20 KO-het perturbations with the highest number of genes DE at p<0.01 in WT background. Blue and red colors represent enrichment of down- and up-regulated genes, respectively. (B) The analogous heatmap for genes DE in Q140/KO-het vs. Q140.

Importantly, at gene level, our datasets reveal novel transcriptional regulatory relationships between perturbed KO-het genes and specific gene sets of high interest. These include well-known MSN cell-type-specific marker genes (Figure 3A), such as D1-MSN genes (e.g. Pdyn, Tac1, Slc35f3) and D2-MSN genes (e.g. Penk, Gpr6, Gprin3), offering candidate upstream regulators of these genes. The enrichment analysis also reveals KO-het influencing gene expression specific to other striatal cell types such as interneurons (Vrk1) and oligodendrocytes (Figure 2A, 3A). In addition to cell-type-enriched genes, our KO-het datasets also reveal KO-het genes that significantly alter the expression of genes related to specific aspects of neuronal biology, such as neuronal activity-dependent genes (Wang et al., 2022).

**Figure 3.**
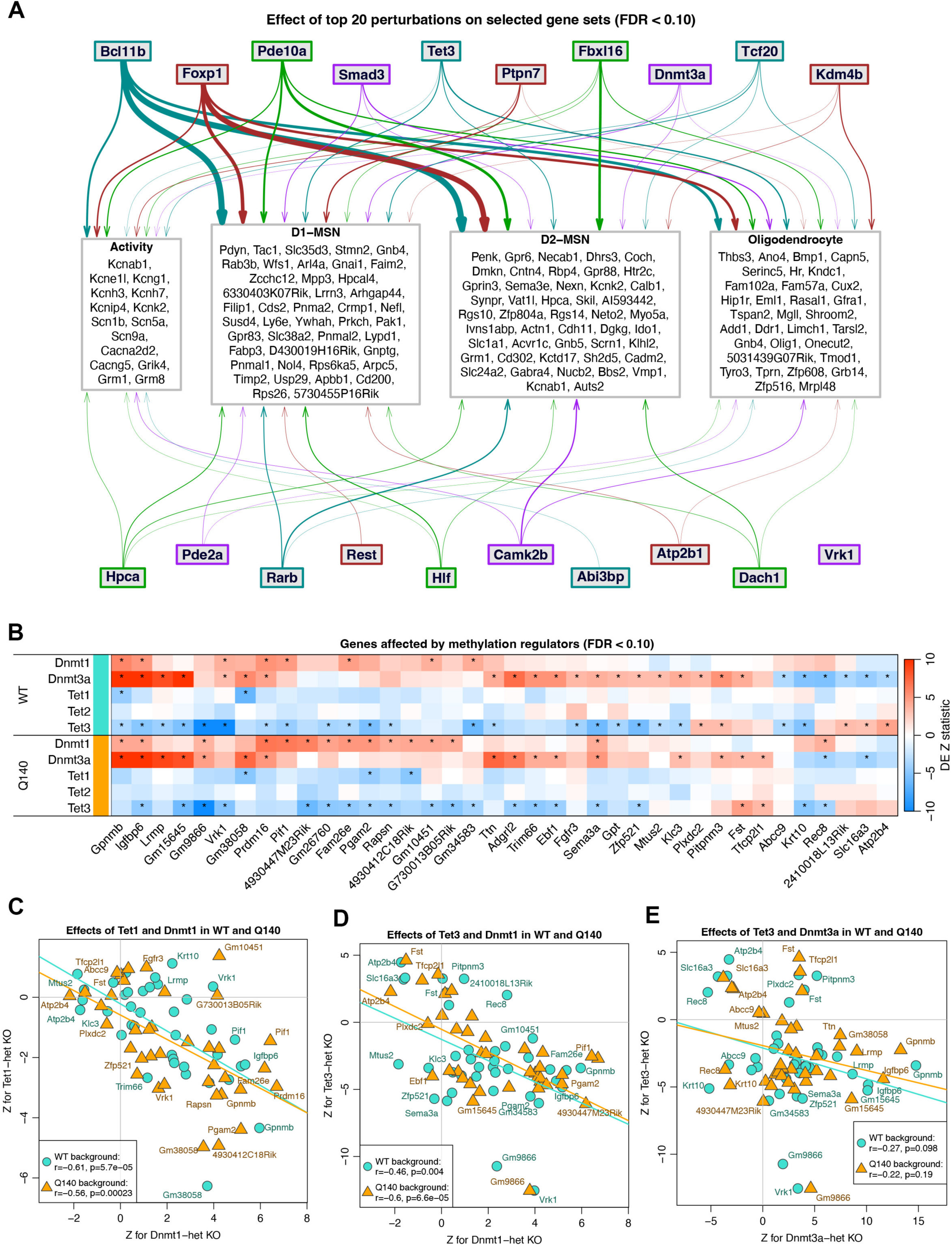
Examples of biological insights from heterozygous knockout perturbations of striatal transcriptomes in wildtype mice. (A) Visualization of effects of the 20 perturbations with highest numbers of genes DE at FDR<0.1 on four categories (activity-dependent genes and markers of D1-MSN, D2-MSN and oligodendrocytes) of genes highly relevant to striatal biology. For each of the four categories, we included genes affected by at least 2 perturbations. Width of lines connecting perturbations to the categories is proportional to the number of genes within the category box that are DE in that KO-het vs. WT at FDR<0.1. For easier tracing of lines between the perturbations and the category boxes, each KO-het carries a distinct but otherwise arbitrary color. Although the Vrk1 KO-het induces the 20th highest number of DE genes, it does not affect any of the gene within these 4 categories. (B) Heatmap of DE Z statistics for KO-het vs. WT and Q140/KO-het vs. Q140 for five KO-het genes whose protein products directly affect DNA methylation. Only those target genes (columns) are shown that, either in WT or Q140 background, pass FDR<0.1 in at least one of the DNA methyltransferases (Dnmt1 and Dnmt3a) and in at least one of the Tet genes. Stars indicate tests with FDR<0.1. (C)-(E) Scatterplots of DE Z statistics for selected pairs of Dnmt1 or Dnmt3a and Tet1 or Tet3 KO-hets. These plots only show the genes displayed in panel (B), namely those that are DE at FDR<0.1 in at least one Dnmt and at least one Tet KO-het in either WT or Q140 background. (C), effects of Tet1 (y-axis) vs. Dnmt1 (x-axis); (D) Tet3 vs. Dnmt1; (E) Tet3 vs. Dnmt3a.

Our study involved two DNA methyltransferases (Dnmt1 and Dnmt3a) and three DNA demethylases (Tet1, Tet2, Tet3). We hypothesized and indeed confirmed that a subset of DEGs downstream of these KO-hets are opposingly regulated by these DNA methylation and demethylation enzymes (Figure 3B). Such effects can also be observed based on correlations of Z statistics for the 38 genes that are significantly (FDR<0.1) affected by at least one of the Dnmt and at least one of the Tet perturbations. These correlations are negative for several perturbation pairs, e.g. Tet1 vs Dnmt1 in WT and in Q140 (Figure 3C), Tet3 vs Dnmt1 in Q140 (Figure 3D), and Tet3 vs Dnmt3a in WT (Figure 3E). Interestingly, one of the 32 genes that are significantly and opposingly regulated by these DNA methylation/demethylation enzymes (Figure 3B) is a Parkinson’s disease risk gene Gpnmb (Diaz-Ortiz et al., 2022). Together, these set of genes constitute potential DNA-methylation sensitive target genes in the adult striatum.

### Pairwise transcriptome-wide concordance analyses of KO-het effects across perturbations

We next examined concordance of KO-het effects among the individual genes in WT and Q140 backgrounds (Supplemental Figure S3A, S3B). To this end, we correlated the Z statistics for KO-het vs. WT across genes common to each pair of comparisons, with higher weight assigned to genes strongly DE in Q140 vs. WT (Methods). Although most correlations are weak (|cor| < 0.2), there are multiple moderate to borderline strong correlations (0.2 < |cor| < 0.6). Clustering of selected perturbations reveals several pairs or small clusters of perturbations anchored by pairs: Gpx-Smad3 (cor = 0.4; other perturbations include Cep164 and Aff4), Foxp1-Camk2b (cor = 0.39), Inf2-Fbxl16 (cor = 0.39; cluster includes Htr2c), Ptpn7-Anks1b (cor = 0.41), Syndig1l- Dnmt1 (cor = 0.36; cluster also includes Hpca) and Tbc1d8-Atf6 (cor = 0.35, cluster also includes Atp2b1 and Tet1). The Inf2-Fbxl16-Htr2c cluster is anti-correlated with the Gpx-Smad3-Cep164- Aff4 cluster, with the strongest correlation of −0.47 between Htr2c and Smad3. Furthermore, the Syndig1l-Dnmt1-Hpca cluster is anti-correlated with the Tbc1d8-Atf6-Atp2b1-Tet1 cluster, with the strongest correlation of −0.46 between Hpca and Tbc1d8.

In Q140 backgrounds, the KO-het perturbations can be clustered into two major clusters that are overall anti-correlated (Supplemental Figure S3B). One cluster containing genes including Kcnh4, Atf6, Rest, Itpr1, Slmap, Nfe2l3, and the other cluster containing genes including Scn4b, Dpysl5, Sp9, Arpp19, Hipk4, Camk2b, Hpca, etc. The gene pairs with the highest positive correlation are Kcnh4-Atf6 (cor = 0.53), and the pair with the highest negative correlation is Scn4b-Kcnh4 (cor=-0.62). Such transcriptome-wide correlation and KO-het gene clusters help identify KO genes with common or opposing effects using exclusively our experimental transcriptomic datasets.

### Htt allelic series WGNCA modules are highly preserved and show striatal cell-type-specific gene enrichment

One of the principal goals of this study is to examine how our selected KO-het genes affect the 13 AS CAG-length dependent coexpression modules (Langfelder et al., 2016). Since our current study has generated 903 WT and 900 Q140 striatal RNA-seq datasets, it allows us to more precise DE analyze DEGs in Q140 vs WT striatum at 6m and examine the preservation of the mHtt-associated AS modules. With meta-average analyses of 8 HD vs 8 WT controls - the sample sizes used for each individual KO-het perturbations - we found 4807 genes with FDR<0.1 (Supplemental Table S5). With a cutoff for |Log2FC| > 0.2, we found 899 significant DEGs (Supplemental Table S5). The latter gene list can be considered a gold standard DEGs for Q140 vs WT in striatum at 6m of age.

This dataset also allows us to answer what mHtt CAG-length dependent modules are preserved in this vast Q140 vs WT datasets. Using our established module preservation analysis (Langfelder et al., 2011) we found all the 5 mHtt CAG-length downregulated modules and 8 CAG- upregulated modules are preserved with Z_summary_ > 5 (Supplemental Figure S4A). The modules with the highest module preservation Zsummary scores greater than 20 are M2, M20 and M39.

We next assessed what striatal cell-type-specific marker genes are enriched in these AS mHtt modules using striatal cell-type marker genes from single-nuclei RNA-seq study of WT striatum (Supplemental Figure S4B; Gokce et al, 2016; Saunders et al., 2018). We found the top CAG-downregulated M2 from Langfelder et al. (2016) is highly enriched with D1- and D2-MSN genes, M25 with D2-MSN genes, M11 with oligodendrocyte and astrocyte genes, and M52 with astrocyte and endothelial cell genes. Among the CAG-upregulated modules, M7 is enriched with oligodendrocyte genes and M9 with D1-MSN genes, M9 with D2-MSN genes, and M20 with neural stem cell genes.

Importantly, using our recently generated striatal cell-type-specific DEGs between Q140 vs WT using snRNA-seq data (Wang et al., 2025), we found M2 and M25 are highly enriched with Q140-downregulated genes in both D1- and D2-MSNs, M11 is enriched with downregulated genes in oligodendrocytes; and all of the upregulated modules except M7 are enriched with DEGs upregulated in Q140 in both D1- and D2-MSNs (Supplemental Figure S5B). M7 is enriched with those upregulated in Q140 in oligodendrocytes. In summary, except for the two modules enriched with oligodendrocyte genes (M7 and M11), all strongly CAG-dependent AS modules appear to reflect MSN biology and specifically either downregulation of their identity genes (M2 and M25) or aberrant gene upregulation (M1, M9, M10, M20, M39, M43, M46).

### Assessing at striatal transcriptome-wide and CAG-associated WGNCA module levels the effects of hemizygosity perturbations in wildtype mice

Since each KO-het perturbation was performed in both WT and Q140 backgrounds, we first asked which perturbations in WT background exert striatal transcriptome-wide or module level effects that mimic those of Q140.

For transcriptome-wide analysis, we used our recently developed transcriptome-wide Z statistics correlation, asking whether Z statistics of KO-het vs WT is significantly correlated with those of Q140 vs WT (Wang et al., 2025). Estimating significance of transcriptome-wide correlations of DE statistics is challenging because of the strong but difficult to estimate gene-gene correlations. Using our large number Q140 and WT samples, we carried out a study of correlations of Z statistics that result from DE analysis between biologically equivalent groups (i.e., DE analysis of null perturbations). This analysis suggests that transcriptome-wide reversal or exacerbation correlations with absolute values below 0.2 should be considered non-significant (p ≥ 0.05, see Methods). We found that Z statistics for KO-het of five genes (Foxp1, Camk2b, Bcl11b, CD4, and Atp2b1) show positive correlations with Q140 vs WT above 0.2 (Figure 4). Among these genes, two of them (Foxp1, Bcl11b) are considered MSN identity genes and are known to drive MSN-specific gene expression (Anderson et al., 2020; Arlotta et al., 2008), and Bcl11b KO has been shown to mimic HD transcriptomic signature (Song et al., 2022). Impressively, Foxp1 KO-het shared 395 downregulated genes and 216 upregulated genes with Q140 vs WT and showed an overall transcriptome-wide Z statistics correlation of 0.31 (Figure 4B). Intriguingly, in WT background, there is only one KO-het (Srebf2) that altered the striatal transcriptome in the direction that is anti-correlated with the effects of Q140 vs WT and passed the threshold |cor| > 0.2. Srebf2 is a M2 module gene that significantly downregulated in Q140 vs WT (Supplemental Table S5), and is found to be at a HD GWAS locus modifying age at onset (Genetic Modifiers of Huntington’s Disease, 2025). However, more studies are needed to fully examine whether Srebf2 genetic manipulations can alter the disease related phenotypes in Q140.

**Figure 4.**
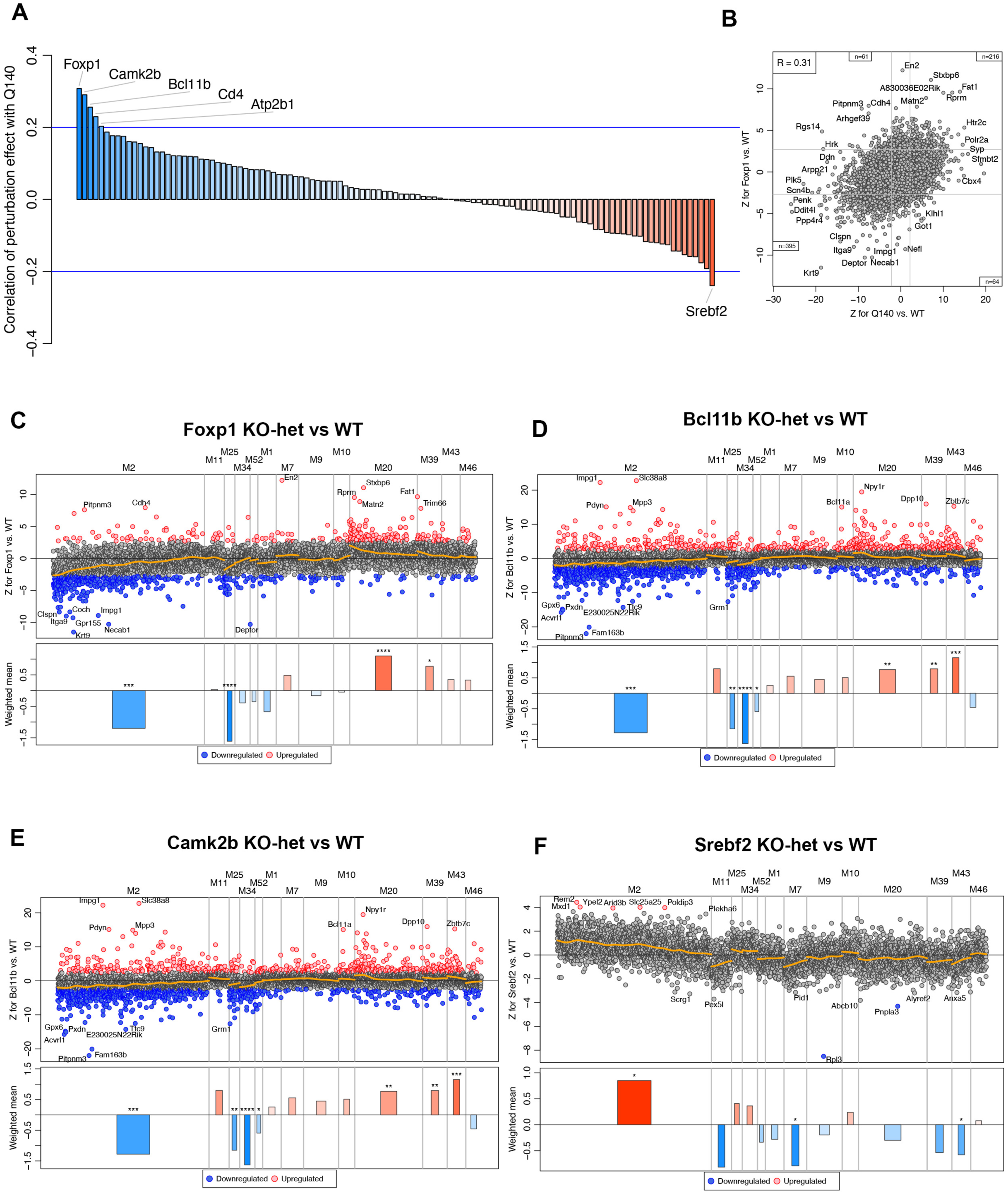
Effects of heterozygous knockout perturbations on HD-associated transcriptomes and CAG-dependent modules in wildtype mice. (A) Barplot of transcriptome-wide correlations of Z for KO-het vs. WT with Z for Q140 vs. WT in the common analysis of all Q140 and WT samples. Blue lines indicate the threshold of ±0.2. (B) Transcriptome-wide scatterplot of Z for Foxp1 KO-het vs. WT against Z for Q140 vs. WT in our common analysis of all Q140 and WT samples. Grey lines indicate approximate locations of FDR=0.1 thresholds. Insets indicate numbers of genes in each of the corner regions (FDR<0.1 in both DE tests) of the plot. Correlation (R) is shown in upper left corner. (C)-(F) Manhattan plots of the effect of KO-het vs. WT for Foxp1 (C), Bcl11b (D), Camk2b (E) and Srebf2 (F). In each panel, the scatterplot on top shows the DE Z statistics for KO-het vs. WT for genes in each of the 13 strongly CAG-dependent modules in Allelic Series striatum. Within each module, genes are ordered from left to right by decreasing module membership *Z.kME*. The orange line shows a local weighted average. A barplot below each scatterplot shows weighted averages of the gene statistics; stars indicate nominal significance (*: p<0.05, **: p<0.01, ***: p<0.001, ****: p<0.0001).

To evaluate at finer resolution the impact of individual KO-het on the WGCNA modules originally defined as Htt CAG-length associated (Langfelder et al., 2016), we created a new plot called WGCNA Module “Manhattan Plot” (see Methods). Briefly, genes in each of the 13 striatal Htt CAG-associated modules are shown along x-axis in order of their module membership as measured by the Z statistic of their correlations with module eigengenes (in WGCNA notation, Z of kME in own module). The y-axis shows the DE Z statistics of the module genes for KO-het vs WT; significantly (FDR<0.1) down- and upregulated genes are shown in blue and red colors, respectively. A local average of the Z statistics is shown as a yellow line; this indicates the overall trend of the KO-het effect. Analogous plots can also be constructed for DE in the Q140 background. We find it informative to use the Manhattan Plot layout to show the reversal score (Methods) instead than DE Z statistics for each gene. In this reversal Manhattan Plot, a gene is shown respectively in blue or ref color if it is significantly reversed or exacerbated, i.e., two conditions are met: (i) It is significantly DE in Q140 vs. WT; and (ii) it is also significantly DE in Q140/KO-het vs Q140 and its expression moves towards (reversal) or away from (exacerbation) the WT expression value.

For each module and KO-het, we summarize the individual gene DE Z statistics or reversal scores using a weighted mean in which hub genes have higher weights than peripheral genes (Methods), resulting in a single measure of the perturbation effect (DE or reversal/exacerbation) on the module. Using our null perturbation analysis, we are also able to create sampling-based significance estimates for each module KO effect or module reversal. Taken together, the module Manhattan plot provides a more nuanced view on how each perturbation - in either the WT or Q140 backgrounds - alters the striatal transcriptome known to be altered by mHtt at the level of gene modules.

Module Manhattan plots for FoxP1 and Bcl11b KO-het (Figure 4C-4F) indicate these perturbations downregulate genes in several mHtt CAG-downregulated modules (M2, M25, M34), while also upregulating genes in the CAG-upregulated modules (M20, M39, and M43). Interestingly, Sreb2 KO-het exhibits the opposite transcriptomic effects compared to Q140 on both CAG-down (M2) and up-regulated modules (M7, M43). These module-level analyses provide a detailed view on the transcriptomic similarity between the KO-het perturbed transcriptomes and that of Q140 at the resolution of CAG modules as well as module hub genes.

### Assessing the effects of hemizygosity perturbations in striatum of HD mice at transcriptome-wide as well as at the level of CAG-associated WGNCA modules

We next address which KO-het can significantly modify Q140 dysregulated striatal transcriptome and AS CAG-modules. We find four perturbations (Scn4b, Sp9, Hpca, Morf4l1) that exacerbate the Q140 transcriptomes with transcriptome-wide reversal Z correlations below the threshold of −0.2 and another four perturbations (Pdp1, Kcnh4, Tet3, and Cpox) that show the opposite, reversing effects above 0.2 (Figure 5A). Among these, perturbations of two genes stand out for showing among the most robust correlation effects compared to Q140 vs WT, Scn4b (cor = −0.42) and Kcnh4 (cor = 0.33); and the other gene with the most robust rescue reveral effect is Pdp1 (cor = 0.38). The other five perturbations mentioned above show more modest reversal correlations close to the threshold of ±0.2.

**Figure 5.**
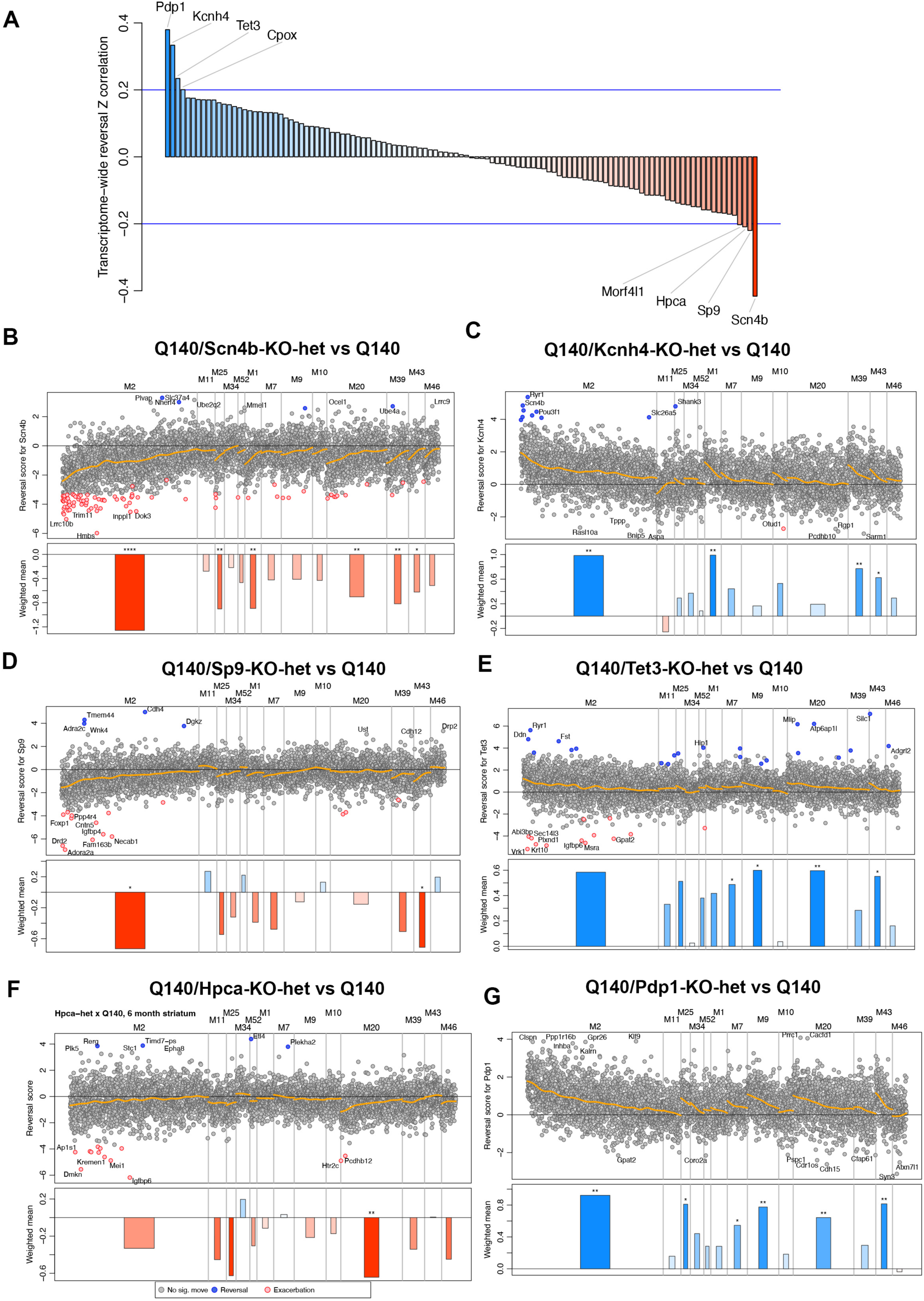
Effects of heterozygous knockout perturbations on HD-associated transcriptomes and CAG-dependent modules in Q140 HD knockin mice. (A) Barplot of transcriptome-wide reversal correlations of Z statistics, i.e., scatterplots of Z for Q140 vs. Q140/KO-het against Z for Q140 vs. WT in the common analysis of all Q140 and WT samples. Blue lines indicate the threshold of ±0.2. (B)-(G). Reversal Manhattan plots for various Q140/KO-hets, i.e., plots of the reversal scores (Methods) for Scn4b (B), Kcnh4 (C), Sp9 (D), Tet3 (E), Hpca (F), Pdp1 (G). In each panel, the scatterplot on top shows the DE Z statistics for KO-het vs. WT for genes in each of the 13 strongly CAG-dependent modules in Allelic Series striatum. Within each module, genes are ordered from left to right by decreasing module membership Z.kME. The orange line shows a local weighted average. A barplot below each scatterplot shows weighted averages of the gene statistics; stars indicate nominal significance (*: p<0.05, **: p<0.01, ***: p<0.001, ****: p<0.0001).

We next created allelic series WGCNA module reversal Manhattan plots for these perturbations (Figure 5B-5G). Impressively, Scn4b KO-het significantly exacerbates 6 AS WGNCA modules (five of which at p<0.01, Figure 5B), which include downregulating genes in the two CAG-downregulated modules (M2, M25) and upregulating genes in the four CAG- upregulated modules (M20, M1, M43 M52). Importantly, within each module the exacerbation is strongest for the module hub genes, which are located towards the left side of the plot for each module (Figure 5B). The other two KO-het genes (Sp9 and Hpca) that exacerbate Q140 transcriptomes appear to show more module-specific effects. Sp9 is a gene known to promote D2- MSN identity gene expression (Xu et al., 2018). Consistent with this function, Sp9 KO-het in Q140 (Fig. 5D) significantly worsens M2 module genes, especially several known D2-MSN enriched genes (e.g. Drd2, Adora2a, Foxp1, Necab1, Fam163b). Hpca encodes the hippocalcin gene, which regulates calcium signaling and neuronal excitability (Park, 2025). HPCA is known to be mutated in autosomal recessive dystonia (Charlesworth et al., 2015) but its underlying pathogenic mechanism is unknown. In our analysis (Fig. 5F), Hpca KO-het significantly exacerbates M20, suggesting that haploid insufficiency of Hpca alters this module in the same direction as mHtt in AS (Figure 5D), providing several shared molecular signatures (e.g. Htr2c). Manhattan plots of the three KO-het genes (Kcnh4, Tet3, Pdp1) that are reversing the Q140 transcriptome all showed significant rescuing of four modules each (Fig. 5C, 5E, 5G). While they all reverse multiple mHtt upregulated modules (e.g. M7, M20, M39, M43), only Pdp1 and Kcnh4 showed significant amelioration of M2 module genes. Interestingly, Kcnh4 KO-het in Q140 background significantly rescued multiple M2 hub genes, including Scn4b and Ryr1 (Figure 5C).

We next created module reversal scores for all AS CAG-dependent modules and for all perturbations (Supplemental Figure S5A, S5B; Supplemental Table S6). We can rank the exacerbations and reversals at individual module levels, such as the top CAG-downregulated module (M2) and CAG-upregulated module (M20). This ranking allows more detailed evaluation of individual perturbations that significantly affect specific CAG-dependent modules but do not elicit a strong transcriptome-wide perturbation effect.

### Similarities of reversal/exacerbation effects among allelic series CAG modules

We next examine whether module gene expression changes induced on the 13 mHtt CAG-dependent modules by our 115 perturbations show correlations in either WT or Q140 backgrounds (Supplemental Figure S6A, S6B). In WT background, we found two module blocks show positive correlation within their own block and anti-correlation across the blocks, with each block containing modules both up- and downregulated by mHtt CAG length (Langfelder et al., 2016). In Q140 backgrounds, the module correlation structure in response to KO-het is only partially shared with those in WT. Interestingly, in both backgrounds, perturbation effects on the modules enriched with MSN genes - M2 and M25 - are highly correlated (Cor = 0.7 in WT; 0.68 in Q140), as are the effects on modules enriched with oligodendrocyte genes (M7 and M11, Cor = 0.89 in WT, Cor = 0.77 in Q140) and astrocyte genes (M52). Importantly, the MSN and oligodendrocyte gene perturbation effects belong to opposite correlation groups and are highly anti-correlated in both WT and Q140, e.g. M2 vs M7 (cor = −0.61 in WT; −0.56 in Q140), suggesting potential coordinated neuronal-glial transcriptional response of these cell types to KO-het perturbagens.

An enigma among the AS modules is M20. It is the top CAG-upregulated WGCNA module and is strongly preserved in expression data from this study, but its enrichment in literature gene sets is relatively weak, suggesting it contains upregulated genes that are somewhat specific to the mHtt effects. Our analysis of module perturbations shows that effects on M20 are most anti-correlated with those on M25 (cor=−0.68 in WT, −0.74 in Q140). Since M25 is enriched with MSN identity genes as well as synaptic genes (Supplemental Figure S4B; Langfelder et al., 2026), this finding suggests that perturbation that upregulate M20 module genes are also likely to downregulate MSN-specific genes and synaptic genes.

### Testing whether module hub genes tend to alter genes within its own module

The WGCNA module hub genes are the most connected gene nodes within a coexpression module (Langfelder and Horvath, 2008), and have been found to be functionally important at least in some biological contexts, e.g. cancer driver genes tend to be module hub genes (Langfelder et al., 2013). We sought to address whether perturbations of hub genes within an AS module tends to alter the genes in the same module compared to other modules. To gain statistical power, we combined the perturbations in both WT and Q140 background and used all the perturbations that have a Z association of at least P<0.1 in a given module (Supplemental Figure S7A, S7B). This analysis reveals that module hub status for M2 is significantly associated with changing the expression of genes in M2 module (cor=0.39, p=0.036). However, our study did not find hub gene status for M20 is significantly correlated with transcriptomic effects in M20, but this could be due to relatively few M20 genes being perturbed in our study. Overall, our study provides yet another example that WGCNA module hub genes could be more central to the biology of genes represented in the module.

### FoxP1 heterozygous knockout exacerbates striatal pathologies in Q140 mice

Foxp1 is one of our top KO-het perturbations -in either WT or Q140 backgrounds - that elicit HD-like striatal transcriptomes, as it shares >600 significantly up- and down-regulated genes with DE in Q140 vs. WT (Figure 4B). Therefore, we examined in more detail the molecular and pathological impacts of Foxp1 KO-het in vivo. We first found both Q140 and Foxp1 KO-het reduces Foxp1 transcript levels by 15-20%, while Q140/Foxp1 KO-het further significantly reduces the transcript to about 65% of WT levels (Figure 6A). Enrichment analysis confirms highly significant enrichment of both up and down regulated genes in both Foxp1 KO-het vs WT as well as Q140/Foxp1 KO-het vs Q140 with those of allelic series KI gene signatures (Figure 6B), and downregulated genes also are enriched with striatal marker genes, D2-MSN marker genes, as well as genes involved in Parkinson’s disease, Alzheimer’s disease, and oxidative phosphorylation.

**Figure 6.**
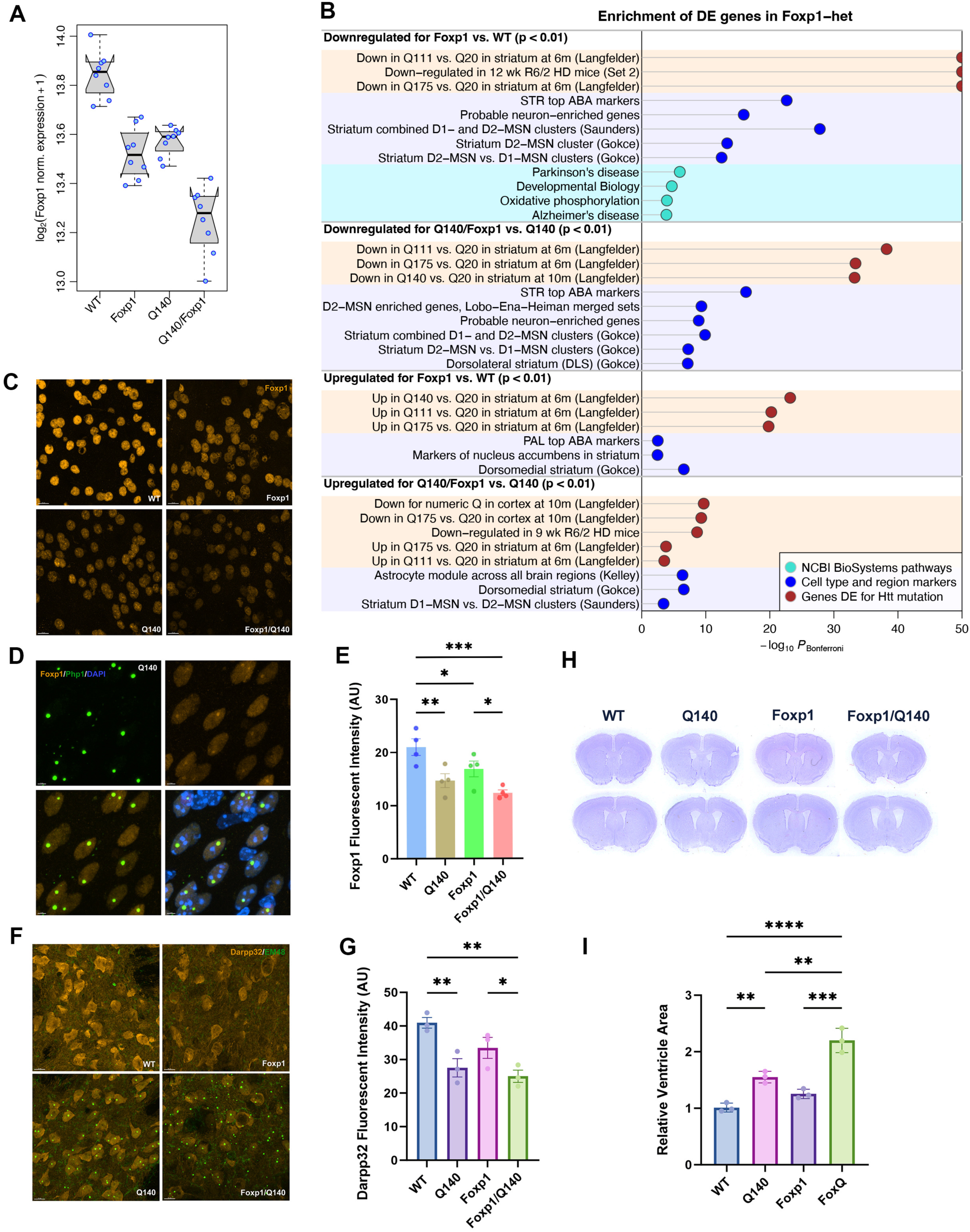
Foxp1 heterozygous knockout exacerbates HD-associated striatal pathology. (A) Boxplot of log-transformed normalized expression of Foxp1 in the 4 genotypes. Blue points show individual expression values, boxes show the median and interquartile range, notches indicate median confidence interval. (B) Enrichment of genes DE in Foxp1 KO-het vs. WT and Q140/Foxp1 KO-het vs. Q140 in selected terms that include BioSystems pathways, cell type markers and genes DE in studies of HD models. For clarity, the enrichment p-values are limited to 10-50. (C) Representative images of Foxp1 staining in striata of 6m Q140/Foxp1 KO-het cohort. Coronal striatal sections (30µm thickness). (D) Foxp1 co-localized with nuclear inclusions in the striatum of Q140 mice. (E) Foxp1 intensity in striata of 6m Q140/Foxp1 KO-het cohort (n=4 per genotype). For all bar graphs in this figure, data are represented as mean ± SEM. One-way ANOVA with Tukey’s HSD tests: * p<0.05, ** p<0.01, *** p<0.001, ****p<0.0001. (F) Representative images of EM48-stained mHtt aggregates in striata of 6m Q140/Foxp1 KO-het cohort. Coronal striatal sections (30µm thickness) were double stained with anti-Darpp32 (amber) and anti-mHtt (EM48, green) antibodies. (G) Darpp32 intensity in striata of 6m Q140/Foxp1 KO-het cohort (n=3 per genotype). (H) Representative Nissl-stained mouse brain coronal sections among 4 genotypes of 6m Q140/Foxp1 KO-het cohort (Bregma 0.65mm and 0.10mm). (I) Quantification of relative ventricle areas in 6m Q140/Foxp1 KO-het cohort (n=3 per genotype).

We examined the neuropathological phenotypes in FoxP1 KO-het alleles in WT and Q140 backgrounds. We found a significant reduction of Foxp1 immunostaining signals in the striatal MSNs in Q140 as well as Q140/Foxp1 KO-het compared to WT, with Q140/Foxp1 KO-het further reduced the Foxp1 protein levels compared to the other genotypes (Figure 6C, 6E). We next examined additional neuropathological phenotypes in HD mice that include mHTT aggregates stained with PHP1 antibody as well as the expression levels of Darpp32, an MSN marker gene (Wang et al., 2025). We found that, compared to Q140, Q140/Foxp1 KO-het have similar levels of mHtt aggregation but significant downregulation of Darpp-32 expression (Figure 6F, 6G). The latter represents an exacerbation of this MSN marker gene dysregulation that is already present in Q140. Interestingly, we found FoxP1 co-localizes with mHtt nuclear inclusions in the striatal MSNs of Q140 mice (Figure 6D), and such Foxp1 pathology was not found in WT striatal MSNs. This finding is consistent with the interpretation that FoxP1 and its close paralog FoxP2, both transcription factors with a long polyglutamine domain, are prone to co-assembly with other nuclear polyQ proteins including mHTT (Hachigian et al., 2017; Saad et al., 2025). Such pathological polyQ co-assembly provides yet another potential mechanism for mHtt to induce Foxp1 loss-of-function beyond its level reduction in HD MSNs.

We further examined ventricular size, as its reduction has been used as indication of forebrain atrophy in both HD mouse models and patients (Hobbs et al., 2010; Marangoni et al., 2014; Zhang et al., 2010). We found 6m old Q140 mice showed a significant increase in ventricular areas compared to WT littermates, while Q140/Foxp1 KO-het showed further significant increase compared to both WT and Q140 (Figure 6H, 6I). These findings support the role of Foxp1 as a rate-limiting protective factor against mHtt-induced toxicity, and its deficiency led to HD-like transcriptome in WT mice and exacerbate forebrain atrophy pathology in HD mice.

### Opposing Neuropathological impacts of Scn4b and Kcnh4 in HD mouse models and patient-derived MSNs

One important discovery in our study is that the two genes with striatum-enriched expression and opposing function (Scn4b and Kcnh4) appear to opposingly regulate mHtt dysregulated transcriptome in HD mice. Scn4b is a modulatory subunit of voltage-gated sodium channel with expression highly specific to the MSNs and is known to modulate MSN excitability (Miyazaki et al., 2014). Meanwhile, Kcnh4 is a poorly studied voltage-gated potassium channel with enriched expression in the striatum and prefrontal cortex (https://www.mousephenotype.org/). Although Kcnh4 has never been studied in the mouse brain in vivo, channels of this type often function in a manner opposing to the voltage-gated sodium channel to regulate neuronal excitability. Given the known increased MSN excitability in HD mice (Veldman and Yang, 2008; Beaumont et al., 2006), and our finding that perturbing these channels appear to dichotomously alter the HD striatal transcriptome, our findings suggest a close relationship between genes regulating MSN excitability and transcriptomic dysregulation of the latter.

We next examined in more detail three genes with most significant impacts on exacerbating (Scn4b) or reversing (Kcnh4) of HD transcriptomes in Q140 (Figure 7A, 7B). Based on transcriptome-wide Z statistics correlation analysis, Q140/Scn4b KO-het vs Q140 has the most negative correlation compared to Q140 vs WT (Cor = −0.42), while Q140/Kcnh4 KO-het vs Q140 shows the highest positive correlation (cor = 0.33). Impressively, if we consider Allelic Series KI lines other than Q140 as perturbations over a Q140 background (by carrying out a DE analysis between each line and Q140 and comparing that to the reference Q140 vs WT), we found that the degree of transcriptomic exacerbation by Scn4b KO-het is comparable to - or even slightly worse than - that of Q175, a mouse model with an inherited ∼190 CAG repeats in mHtt (Menalled et al., 2012) (Figure 7C). Neuropathologically, we found the striatum of Q140/Scn4b KO-het at 6m of age showed significant increase in both PHP1^+^ mHtt aggregation and decrease in Darpp32 MSN marker protein expression (Figure 7D-7F).

**Figure 7.**
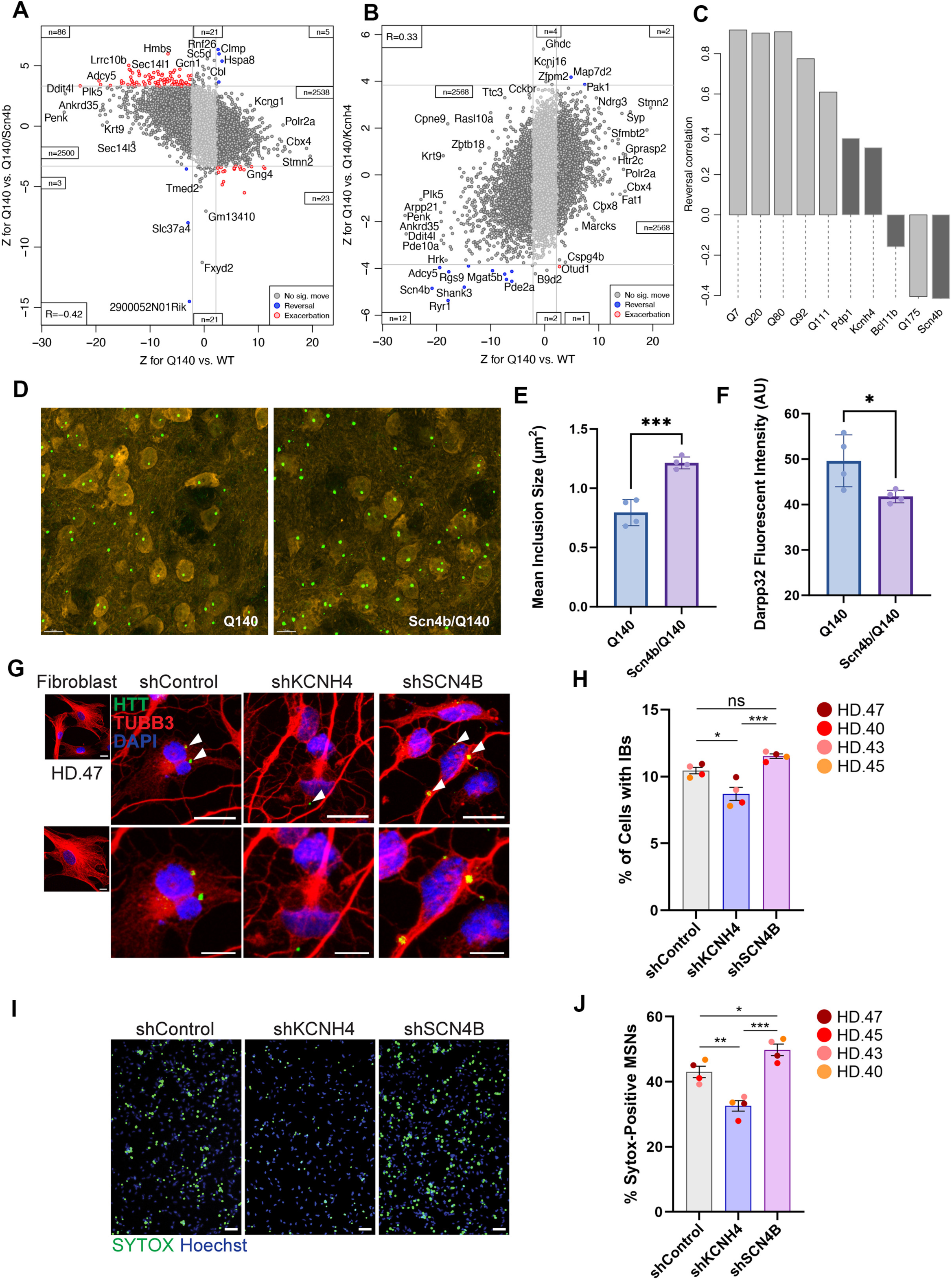
Effects of Scn4b and Knch4 perturbations in HD-associated phenotypes in HD mouse model and patient-derived MSNs. (A), (B) Z statistics reversal scatterplots for Q140/Scn4b KO-het and Q140/Kcnh4 KO-het. In each plot, x- and y-axes show Z for Q140 vs. WT in our analysis of all Q140 and WT samples and Z for Q140 vs. Q140/Scn4b KO-het (A) and Q140 vs. Q140/Kcnh4 KO-het (B), respectively. Grey lines indicate approximate locations of FDR=0.1 threshold. Blue and red color indicate genes reversed and exacerbated, respectively, at FDR<0.1. Insets indicate numbers of genes in corner regions (DE at FDR<0.1 in both tests). Transcriptome-wide correlation R is shown in a separate inset. (B) Transcriptome-wide Z statistics reversal correlation (Methods) for Q lengths other than Q140 in the allelic series (Langfelder et al., 2016) considered a perturbation of Q140 and four other perturbations, namely Q140/Kcnh4 KO-het,, Q140/Pdp1 KO-het, Q140/Bcl11b KO-het and Q140/Scn4b KO-het. (C) Representative images of EM48-stained mHtt aggregates in striata of 6m Q140 and Scn4b/Q140. Coronal striatal sections (30µm thickness) were double stained with anti-Darpp32 (amber) and anti- mHtt (EM48, green) antibodies. (D) Average sizes of nuclear inclusion (NI) in 6m Q140 and Q140/Scn4b KO-het striata (n= 4 per genotype). For all bar graphs in this figure, data are represented as mean ± SEM. Student’s t-test, *** p<0.001. (E) Darpp32 intensities in 6m Q140 and Q140/Scn4b KO-het striata (n= 4 per genotype). Student’s t- test, *p<0.05. (G and H) Representative images (G) and quantification (H) of cells with HTT inclusion bodies (IBs) from four independent HD-MSNs (n = 4) transduced with shControl, shKCNH4, or shSCN4B. Cells were immunostained with anti-mHTT (MW8) and TUBB3 antibodies. An average of 277 cells per sample was counted in three or more randomly chosen fields. Scale bars represent 20 μm (top) and 10 μm (bottom). (I and J) Representative images (I) and quantification (J) of SYTOX-positive cells as a fraction of Hoechst-positive cells in HD-MSNs derived from four independent HD individuals (n = 4) transduced with shControl, shKCNH4, or shSCN4B. Scale bars represent 100 μM. Statistical significance was determined by one-way ANOVA with Tukey’s post hoc test. **p<0.01, *p<0.05, ns, not significant. The sample size (n) corresponds to the number of biologically independent samples; GM04198 (CAG repeat size 47; HD.47), ND33947 (HD.40), ND30013 (HD.43), and GM04230 (HD.45). Each dot represents one individual’s reprogrammed HD-MSN.

To further validate this finding in HD patient neuronal models, we used patient fibroblast-derived directly reprogrammed MSNs that have been shown to recapitulate both inclusion body formation and HD disease-stage-dependent MSN loss (Victor et al., 2018; Oh et al., 2022). We used a scrambled control shRNA or shRNAs against SCN4B or KCNH4 to knockdown (KD) these genes in directly reprogrammed MSNs derived from multiple HD patients (HD-MSNs) (Figure 7G to 7J). We found that KCNH4 KD significantly reduced mHTT inclusion bodies (IBs) in HD-MSNs compared to control shRNA, while SCN4B KD showed a trend of increasing IBs (Figure 7G, 7H). Importantly, SCN4B KD significantly exacerbated neuronal death in HD-MSNs whereas KCNH4 KD significantly lowered neuronal death in HD-MSNs (Figure 7I, 7J). Together, these results demonstrate that, in both HD mouse models and human HD patient-derived MSNs, the two ion channels with selectively enriched expression in MSNs, Scn4b and Kcnh4, are likely to play important opposing roles in striatal MSN-selective pathogenesis in HD and hence should be further explored for potential therapeutics.

### Evidence for metabolic enzyme Pdp1 in modifying MSN pathogenesis in female HD mice

Pyruvate dehydrogenase phosphatase 1 (Pdp1) is an important metabolic enzyme in mitochondria that regulates aerobic metabolism by controlling the activity of the pyruvate dehydrogenase complex (PDC; Kumar & Greenberg, 2025). PDC is essential to convert pyruvate to acetyl-CoA, an essential substrate for ATP production in mitochondria through the Tricarboxylic Acid cycle (TCA) cycle (Kumar & Greenberg, 2025). While mutant and wildtype HTT are known to play a role in influencing ATP/ADP ratio in cells and mitochondria oxidative phosphorylation (Ismailoglu et al., 2014; Seong et al., 2005), how it may interact with key metabolic enzymes and modify HD pathogenesis remains unclear.

Pdp1 transcripts are significantly downregulated in Q140 striatum, and its levels are further significantly downregulated in Q140/Pdp1 KO-het (Figure 8A). Although Pdp1 perturbation in Q140 background did not induce any significant DEGs in WT or Q140 background (FDR < 0.1), we found that the transcriptome-wide Z reversal statistics ranked Pdp1 as the top KO-het perturbation in reversing the dysregulated Q140 striatal transcriptome compared to that of WT (Fig. 5A; Fig. 8B). And such significant reversal effect by Pdp1 KO-het is also evident in the analyses of CAG-dependent modules (Fig. 5G). Interestingly, transcriptome-wide Z statistics reversal effects appear to be more robust in the female samples compared to the male samples (Figure S8A, S8B), but more studies are needed to confirm such a sex-dependent effect. Enrichment analysis - using DEGs between Q140/Pdp1 KO-het vs Q140 at a lower statistical threshold (i.e. p < 0.01) - reveals a significant reversal of the downregulated genes in in the HD mice; and also upregulate the MSN identity genes (Figure 8B, 8C).

**Figure 8.**
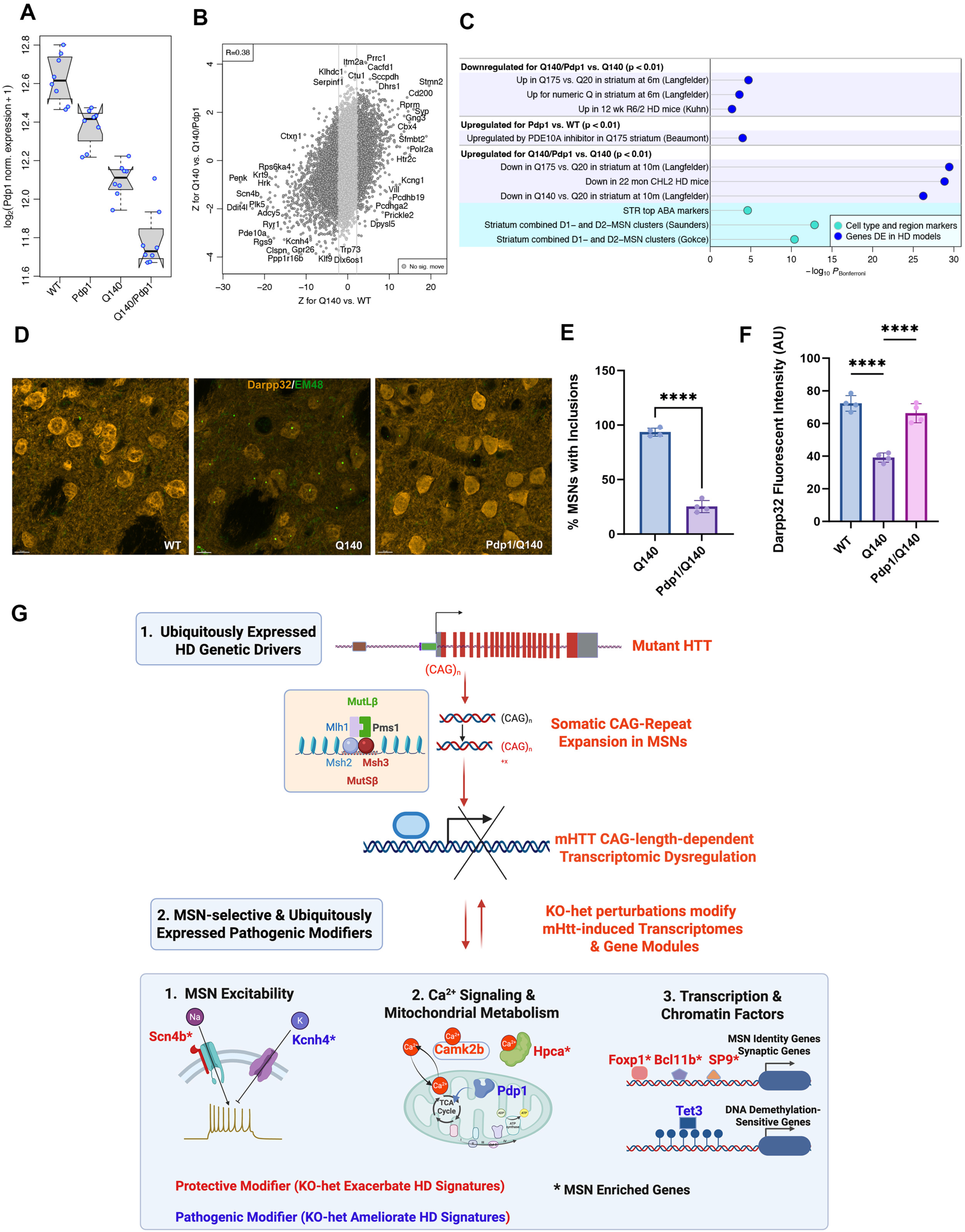
Pdp1 heterozygous knockout effects on striatal neuronal pathology and summary schematics on novel modifiers of striatal transcriptomic networks and neuronal pathology in vivo. (A) Boxplot of log-transformed normalized expression of Pdp1 in the 4 genotypes. Blue points show individual expression values, boxes show the median, median confidence interval, and interquartile range. (B) Reversal scatterplot for Q140/Pdp1 KO-het. X-axis shows Z for Q140 vs. WT in our analysis of all Q140 and WT samples, while the y-axis shows Z for Q140 vs. Q140/Pdp1 KO-het. Grey lines indicate approximate locations of FDR=0.1 threshold. Blue and red color indicate genes reversed and exacerbated, respectively, at FDR<0.1. Insets indicate numbers of genes in corner regions (DE at FDR<0.1 in both tests). Transcriptome-wide correlation R is shown in a separate inset. (C) Enrichment of genes DE in Q140/Pdp1 KO-het vs. Q140 in selected terms that include cell type markers and genes DE in studies of HD models. For clarity, the enrichment p-values are limited to 10-50. (D) Representative images of EM48-stained mHtt aggregates in striata of 6m Q140/Pdp1 KO-het cohort. Coronal striatal sections (30µm thickness) were double stained with anti-Darpp32 (amber) and anti-mHtt (EM48, green) antibodies. (E) Percentage of medium spiny neurons (MSNs) with NIs in striata of 6m Q140 and Q140/Pdp1 KO-het (n= 4 per genotype). For both bar graphs in this figure, data are represented as mean ± SEM. Student’s t- test, **** p<0.0001. (F) Darpp32 intensity in striata of 6m Q140/Pdp1 KO-het cohort (n= 4 per genotype). One-way ANOVA with Tukey’s HSD tests: **** p<0.0001. (G) The top genes and their implicated biological pathways revealed by this large-scale perturbation of mHtt CAG-dependent module hub genes are illustrated here. MSN-selective transcriptionopathy is downstream of somatic mHtt CAG repeat expansion driven by MMR genes (particularly Msh3 and Pms1, Wang et al., 2025). Genes dysregulated by mHtt – some are selectively expressed in striatal MSNs and others are more broadly expressed - can further exert impacts on MSN-selective pathogenesis in HD mice.

We next examined whether the transcriptomic effects of Pdp1 KO-het in Q140 could be reflected in modification of HD-related neuropathology. Compared to Q140 littermates, Q140/Pdp1 KO-het mice show a significant reduction of mHtt aggregation load (Figure 8D, 8E), and significant upregulation of the Darpp32 immunostaining in the striatum (Figure 8D, 8F). Together, this evidence suggests Pdp1 downregulation is a protective response that can ameliorate certain transcriptomic and neuropathological changes in Q140 MSNs. Moreover, since Pdp1 reduction reduces PDC activation, acetyl-CoA levels, and likely ATP production via TCA cycle, this result also implicates this essential mitochondrial metabolic pathway in aspects of HD MSN-selective pathogenesis in vivo.

## DISCUSSION

Our study provides an important proof-of-concept that germline heterzygous knockout perturbation with transcriptomic readout is a valuable, scalable approach to evaluate gene function and discover disease modifiers in vivo. We describe a large-scale perturbation of 115 heterozygous knockout alleles in WT and HD mice with striatal RNA-seq as the primary readout. We demonstrate a scalable pipeline that includes the generation of 32 mice per KO allele (WT, KO-het, Q140, Q140/KO-het) and producing a dataset of 32 striatal RNA-seq samples. Thus, our study enables analysis of each perturbation individually and permits integrative analyses of all 115 KO-het/RNA-seq experiments. To this end, we developed standardized breeding, RNA-seq, and data analysis pipelines, along with new bioinformatic tools to compare, contrast, and rank transcriptomic modifier effects based on HD gene signatures at transcriptome-wide and individual WGCNA module levels. Among the modifiers of HD-associated transcriptomic networks are MSN-enriched genes (Scn4b, Kcnh4, Foxp1, Bcl11b, and Sp9) as well as ubiquitously expressed genes (Pdp1, Tet3). These genes are involved in several biological pathways, including MSN excitability, MSN-specific gene transcription, Ca^2+^ signaling, and mitochondrial energy metabolism (Figure 8G). Our prior study demonstrated that striatal transcriptomic dysregulation depends on somatic mHtt-CAG repeat expansion, driven by MMR genes including Msh3-Msh2 (MutSβ) and Mlh1-Pms1 (MutLβ; Wang et al., 2026). Taken together, our findings support a model in which somatic expansion-driven transcriptomic dysregulation can further modify MSN-selective pathogenesis by altering the expression of genes in several molecular pathways (Fig. 8G).

Our study provides evidence supporting the selection of WGCNA module hub genes as a source of genetic regulators of gene modules in relevant biological or disease contexts. Despite over 4000 PubMed papers containing the terms “WGCNA” and “hub gene”, to our knowledge, few, if any, have systematically perturbed these hub genes to examine their effects on modular gene expression and underlying biology. Our study represents a rigorous test of whether Htt CAG-dependent striatal module hub genes can modify HD-related transcriptomes, gene modules, and neuropathology. This analysis revealed six genes in WT background and eight genes in the Q140 background that modify HD-associated transcriptomes and modules in vivo. These genes belong to several broad classes, including those involved in MSN excitability (Scn4b, Kcnh4), MSN-specific transcription factors (Foxp1, Bcl11b, Sp9, Srebf2), epigenome/chromatin function (Tet3, Morf4l1), calcium signaling and homeostasis (Camk2b, Hpca, Atp2b1), and energy metabolism (Pdp1). Crucially, several of these new HD transcriptomic modifier genes are expressed selectively to the striatal MSNs, i.e. Scn4b, Kcnh4, Foxp1, Bcl11b, Sp9, providing novel MSN-enriched modifiers of HD transcriptomes. Our study also shows that perturbations of M2 module hub genes is significantly associated with gene expression changes in this module but not in another module (M20). Because most selected KO-het genes are from the M2 module, our study cannot determine whether perturbing genes belong to other Htt CAG-dependent modules similarly affects expression of genes in their own modules in vivo.

Our study provides a rich database of causal effects between genetic perturbations of 115 genes and resulting gene expression changes in the mammalian striatum. This substantially expands our prior study of 52 KO-het only in WT background (Wang et al., 2022). The resulting compendium of 115 KO-het genes and 6,517 significantly dysregulated genes can be readily searched to identify potential gene regulatory relationships. Users can identify different upstream perturbagens that regulate the same downstream gene or genes with shared biological function or pathways. We envision the latter as particularly powerful - for example, using gene set enrichment analysis (i.e. anRichment) terms to identify perturbated genes that modify a specific biological pathway. As a proof-of-concept, we showed that a novel set of genes, when perturbed, alters the expression of striatal D1-MSN or D2-MSN specific genes, ion channels regulating neuronal activity, and genes opposingly regulated by DNA methylation and demethylation enzymes. The latter include Gpnmb, a PD GWAS risk gene, now shown to be likely regulated by two DNA methylation enzymes (Dnmt3a and Dnmt1) and two demethylation enzymes (Tet1, Tet3). This data mining approach should be of interest to those who seek to study the regulation of specific target genes in our database of KO-het regulated genes, which is well beyond the scope of our original goal of studying HD-related gene regulation. This large-scale experimentally generated, causal gene perturbation transcriptomic dataset constitutes a rich resource for further computational modeling and knowledge extraction.

A key mechanistic insight from our study is that two striatal MSN-enriched voltage-gated channels with opposing function on neuronal excitability, Scn4b and Kcnh4, confer dichotomous effects on striatal transcriptome, mHtt aggregation, and patient-derived HD MSN cell death (Figure 7). Scn4b is highly expressed in MSNs and is localized to their axons, and mice deficient in Scn4b show an impairment in resurgent Na+ current, which is required for maintaining the repetitive firing frequency of MSNs (Miyazaki et al., 2014). Scn4b knockout or knockdown also reduces the MSN action potential threshold and synaptic long-term depression (Ji et al., 2017). In the AS transcriptomic study, we found Scn4b downregulation as the most significant CAG-length dependent gene regulation in our study, suggesting a primary role of its dysregulation in HD. Its KO-het effect size in exacerbating the Q140 transcriptome is comparable to that of Q175. Our findings on Scn4b are consistent with another study (Sathitloetsakun et al. BioRxiv, 2026) showing Scn4b KD in wildtype striatum elicits profound locomotor deficits and HD-associated transcriptomic signatures, and overexpression of Scn4b can ameliorate transcriptional and electrophysiological deficits in HD mice. Since our unbiased large-scale genetic perturbations identified an understudied striatum-enriched voltage-gated potassium channel that may ameliorate HD-related molecular and pathological phenotypes, including patient MSN aggregation and cell death, our study suggests rebalancing the channels related to MSN-excitability could be a therapeutic strategy for HD. Since neuronal channels are readily druggable targets, future studies should explore whether augmenting Scn4b levels or function or inhibiting voltage-gated K+ channels such as Kcnh4 could serve as therapeutic strategies to modify HD pathogenesis.

A key advance in this study is the development of a fully integrated heterozygous KO- transcriptomic perturbation pipeline, which is scalable, information-rich (with large-scale transcriptomic datasets), and exquisitely sensitive in detecting modifier effects. The latter is facilitated by incorporating new computational pipeline with DEG analysis for each KO-het perturbation, as well as transcriptome-wide Z statistics correlations and WGCNA module-level statistics (e.g. “Manhattan plots”). This multi-level approach yields high sensitivity to detect genes that exert haploinsufficiency effects on WT or Q140 transcriptomes. An example of this is Pdp1, which shows few DEGs using traditional analysis but ranks highly by transcriptome-wide reversal Z statistics and module-level analysis. These findings were validated by the rescue of striatal neuronal pathology in Q140/Pdp1 KO-het (Fig. 8D-8E). One potential caveat of the approach is that heterozygous KO may be too mild to reveal a gene’s full potential effects in a biological or disease context; thus, negative results from any KO-het gene should not be overinterpreted. Indeed, there may be value in pursuing full KO perturbations (Wang et al., 2025) or AAV-mediated CRISPR KO (Wertz et al., 2020; Zheng et al., 2024) to evaluate gene function in transcriptomic regulation in vivo. Despite this limitation, given the large number of omics-derived candidate genes or proteins relevant to HD (Shirasaki et al., 2012) and other brain diseases (Hartl et al., 2021; Zhang et al., 2013; Parikshak et al., 2015), our module hub gene-based scalable perturbation strategy may help identify disease-modifying genes for other brain disorders in vivo.

## METHODS

### Materials Availability

Mouse lines generated in this study are in the process to be deposited to public repository for distribution to the scientific community. Before the lines become available to public, limited stock will be available upon request following a completed material transfer agreement.

### Data and Code Availability

We also in the process of creating an online searchable database through https://www.hdinhd.org/, which will host the analyses of individual KO-het perturbations in both WT and Q140 backgrounds. Meanwhile, any information related to the data presented in the study, e.g. required to reanalyze the data reported in this work paper, is available from X. W. Yang upon request.

### Experimental animals for the KO-het perturbation studies

The mice use for current study were housed in the vivarium at the University of California, Los Angeles (Los Angeles, California, U.S.A.). 81 KO mouse lines were imported from public repositories (i.e., JAX, MMRRC, EMMA, RIKEN, NCI and TCP): among them, lines from JAX and MMRRC were imported as live animals, lines from other sources were rederived at Cedars-Sinai Medical Center from frozen sperm or embryos. Total 34 new KO mouse lines were generated by Yang Lab using the CRISPR/Cas9 method at the Jackson Laboratory’s Customer Model Generation Core (30 lines) and at Mouse Biology Program at University of California, Davies (4 lines; Supplemental Table S1). 10 lines are conditional knockout mice (Supplemental Table S1), male mice in these 10 lines were first bred to E2a-Cre female (JAX 003724), heterozygous offspring carrying post-Cre (null) allele were used for further crossing with Q140 KI mice (JAX 027409). Each KO heterozygous was bred with Q140 to generate offspring with 4 different genotypes. For each crossing, pups were weaned and ear tagged at around 3 weeks of age, then genomic DNAs extracted from ear biopsies were used for genotyping PCR. All mice were maintained and bred under standard conditions consistent with National Institutes of Health guidelines and approved by the University of California, Los Angeles Institutional Animal Care and Use Committees. The cages were maintained on a 12:12 light/dark cycle, with food and water ad lib.

### Sample collection and RNA-sequencing

Eight mice in each of the 4 genotypes from heterozygous KO (KO-Het) x Q140 mice between 26 weeks and 27 weeks of age were used for transcriptomic study, thus a total of 32 mice were profiled from each cross (sex matched among each genotype). Striatal tissues were harvested in the afternoon within the same time window and freshly frozen on dry ice. Total RNA was extracted using Trizol and then RNeasy kit (Qiagen). All RNA samples had RIN>8.0, with average RIN≈8.6. Library preparation and RNA sequencing were performed by the UCLA Neuroscience Genomics Core (UNGC) or UCLA Technology Center for Genomics & Bioinformatics (TCGB). Libraries were prepared using the Illumina TruSeq RNA Library Prep Kit v2 and sequenced on an Illumina HiSeq4000 or NovaSeq X Plus sequencer using strand-specific, paired-end, 69-mer sequencing protocol to a minimum read depth of 30 million reads per sample. Reads were aligned to mouse genome mm10 using the STAR aligner with default settings. Read counts for individual genes were obtained using HTSeq. RNAseq data has been deposited within the Gene Expression Omnibus (GEO) repository.

### MicroRNA-mediated MSN reprogramming

Direct neuronal reprogramming of human fibroblasts into MSNs was performed as previously described (Victor et al., 2018; Oh et al., 2022). Human fibroblasts were seeded at 300,000 cells per well into a Costar six-well plate (Corning). On Day 0, cells were transduced with 4 mL of a pooled lentiviral cocktail containing pTight-9/9*-124-BclxL, rt TA, CTIP2, DLX1, DLX2, and MYT1L in the presence of polybrene (8 μg/mL) and centrifuged at 1,000 g for 30 minutes at 37 °C using a swinging bucket rotor. The next day (Day 1), cells were washed with PBS and cultured in fibroblast medium (2 mL per well) with doxycycline (Dox, 1 μg/mL). On Day 3, the medium was replaced with fresh fibroblast medium containing Dox (1 μg/mL) and puromycin (3 μg/mL). On Day 5, cells were replated onto poly-L-Ornithine/laminin/fibronectin-coated glass coverslips pretreated with nitric acid, and cultured in fibroblast medium supplemented with Dox (1 μg/mL). The next day (Day 6), the medium was switched to neuronal medium (ScienCell) containing neuronal growth supplement and penicillin/streptomycin, supplemented with Dox (1 μg/mL), valproic acid (1 mM), dibutyryl cAMP (200 μM), BDNF (10 ng/mL), NT-3 (10 ng/mL), retinoic acid (1 μM), RevitaCell Supplement (Gibco, 1X), and antibiotics (puromycin, 3 μg/mL; blasticidin, 3 μg/mL; geneticin, 300 μg/mL). Dox was replenished every 2 days, and half of the neuronal medium was replaced every 4 days. Blasticidin and geneticin were discontinued after Day 10, and puromycin and RVC were stopped after Day 21.

### Immunostaining analysis in reprogrammed MSNs

Cells were fixed in 4% paraformaldehyde at room temperature (RT) for 20 minutes, washed three times with phosphate-buffered saline (PBS) for 5 minutes each, and permeabilized with 0.2% Triton X-100 for 10 minutes at RT. Cells were then blocked with 5% bovine serum albumin (BSA) and 1% goat serum in PBS for 1 hour at RT, followed by overnight incubation with primary antibodies, rabbit anti-tubulin β III (BioLegend, #802001) and mouse anti-mHTT (MW8) (DSHB), at 4 °C. After three 10-minute washes with PBS, cells were incubated with appropriate secondary antibodies, goat anti-rabbit and mouse IgG conjugated with Alexa-488 or -594 (Invitrogen), for 1 hour at RT. Cells were washed twice with PBS for 5 minutes each, then incubated with DAPI (Sigma-Aldrich, D-9542) for 10 minutes at RT. After two 5-minute washes in PBS, cells were mounted on a slide for imaging. Images were captured using a Leica SP5X white-light laser confocal system with LAS Advanced Fluorescence 2.7.3.9723. For each coverslip, three randomly chosen fields of view were imaged, and all fluorescence channels were overlaid. For quantification, positive cells were counted in each field of view and expressed relative to the total number of DAPI-positive cells.

### SYTOX assay

To assess neuronal cell death, 0.1 µM SYTOX Green nucleic acid stain (Invitrogen, S7020) was added to the culture medium, along with 1 µL/mL Hoechst 33342 (Thermo Scientific, 66249). Cells were incubated for at least 30 minutes at 37 °C before live-cell imaging. Images were acquired using a Leica DMI 4000B inverted microscope with Leica Application Suite (LAS) Advanced Fluorescence.

### Mouse brain tissue histology

Mice were perfused and coronal brain sections were cut at 30 µm with Leica Cryostat and stored at −20°C in cryopreservation solution (30% sucrose, 30% glycerol in 1xPBS) for further processing. Immunostaining and quantification were carried out using published methods(Wang et al., 2025). PHP1 antibody (Millipore MABN2490; 1:2,000 dilution) and EM48 (Millipore MAB5374; 1:350 dilution) was used for mHTT aggregate detection; Darpp32 antibody (Cell Signaling 2306; 1:500 dilution) and Foxp1 antibody (Abcam ab16645; 1:500 dilution) were used for detecting these targets. For ventricle size measurement, equivalent first sections were chosen based on common morphological landmarks, every 6th section thereafter was stained with Nissl. The areas of the ventricles were measured using ImageJ and quantified.

## QUANTIFICATION AND STATISTICAL ANALYSIS

### Selection of perturbation genes

The authors of (Langfelder et al., 2016) analyzed striatum expression data from 3 ages (2, 6 and 10 months) and defined consensus co-expression modules across the 3 ages. They further quantified association with CAG length, module membership and module eigengene association with CAG length at each of the 3 ages and summarized each separate statistic across the three datasets by meta-analysis. Given that the number of genes dysregulated in the striatum of Allelic Series mice (Langfelder et al., 2016) is too large to test perturbations of all of them, we aimed to define a single measure reflecting evidence that a gene may be involved in mHtt Q-length-dependent transcriptomic dysregulation. The measure uses statistics for gene association with CAG length, module membership Z statistics together module eigengene association Z statistics with CAG length, as well as Z statistics derived from causal analysis using Network Edge Orienting (NEO) (Aten et al., 2008). Specifically, for a gene *g*, we define the overall NTPS statistic (called *Z.NTPS*) as a weighted sum of three statistics: (1) the meta-analysis Z for CAG length, denoted *Z.CAG*, (2) a Z-like statistic *Z.module* based on consensus module membership and the meta-analysis Z for module eigengene association with CAG length, and (3) a Z-like statistic *Z.causal* reflecting the number of genes whose expression is predicted by NEO to be causally affected by expression of gene *g*.

*Z.module* for gene *g* is defined as

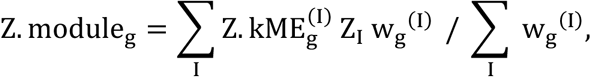

where index *g* labels genes and index *I* labels modules. *Z.kME_g_^(I)^* is the meta-analysis Z statistic for module membership of gene *g* in module *I* (i.e., meta-analysis of the Z statistics corresponding to correlations of the expression profiles of gene *g* with module eigengene of module *I* in the three datasets); *Z_I_* denotes the meta-analysis Z statistic of association of module eigengene of module *I* with CAG length, and *w_g_^(I)^* are weights designed to emphasize modules in which the gene has high kME. We chose the form

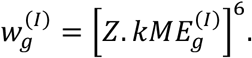

The relatively high power of 6 ensures that the module to which the gene has the highest intramodular connectivity dominates the weighted average.

For the causal analysis using NEO software (Aten et al., 2008), we retained the 6541 genes whose robust correlation with CAG length is at least 0.25 in at least 2 of the 3 striatum age-specific datasets. We then calculated Local Edge Orienting (LEO) scores in each of the 3 striatum sets for all ordered pairs of the 6541 genes. The NEO software outputs, for each putative causal relationship, the LEO score that measures how much better the relationship fits the data than other possible (competing) causal relationships (higher LEO score is better) as well as a p-value for the actual model fit; higher, i.e., less significant p-values indicate a better fit (Aten et al., 2008). For each pair, we summarized the statistics across the 3 datasets using Stouffer meta-analysis Z statistic of the LEO p-values and sum of the LEO scores. We called those edges significant for which the meta-analysis Z statistic of the LEO p-values is less than 2.5, sum of LEO statistics is at least 2, all three individual LEO scores are all above −0.5 and at least two are above 0.5, ensuring a certain level of consistency among the three datasets. For each gene, we counted the number of other genes it is predicted to causally affect based on these thresholds. These counts, referred to as upstream causal connectivities, were then multiplied by the sign of the correlation of the gene expression with CAG length so the signed upstream connectivity is negative for genes downregulated with increasing CAG length. The signed counts were then scaled to unit variance, resulting in a Z-like statistic we call *Z.causal*.

The three Z statistics *Z.CAG*, *Z.module* and *Z.causal* are each divided by its standard deviation, i.e., scaled to variance 1, and the overall *Z.NTPS* statistic is calculated as the weighted average of scaled *Z.CAG*, *Z.module* and *Z.causal* with weights 1, 1 and 0.5, respectively. The lower weight of the causal evidence reflects our lower confidence in the causal analysis than in association- and gene co-expression-based statistics; the causal evidence is largely based on partial correlations that exhibit more variability and susceptibility to noise than marginal correlations.

This list turned out to be dominated by hub genes from Allelic Series module M2, a large (>1800 genes) MSN marker-enriched module whose eigengene was most strongly associated with CAG length in Allelic Series striatum data. Hence, we truncated the list of genes from M2 and added a smaller number of genes from each of the other 12 strongly CAG-dependent modules plus several genes curated manually.

### Bulk RNA-seq data preprocessing

Each cross was analyzed as a separate dataset. In each dataset, we retained only genes with at least 0.06 robust counts per million reads (CPM) in at least the number of samples in the smallest group (genotype with smallest number of samples, generally 8). The rationale for this filtering is to only include genes that are likely to be expressed in at least genotype. The robust CPM is calculated by first normalizing the counts using the Relative Log Expression (RLE) normalization implemented in DESeq2 (Love et al., 2014) and then calculating “counts per million reads” from the normalized data. To identify potential outliers, we used a modified version of the sample network methodology originally described (Oldham et al., 2012). Specifically, to quantify inter-sample connectivity, we first transformed the raw counts using variance stabilization (DESeq2 function varianceStabilizingTransformation which also includes RLE normalization) and then used Euclidean inter-sample distance based on the scaled profiles of the 8000 genes with highest mean expression. The intersample connectivities k were transformed to Z scores using robust standardization,

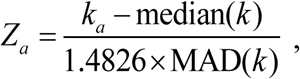

where index a labels samples, MAD is the median absolute deviation, a robust analog of standard deviation, and the constant 1.4826 ensures asymptotic consistency (approximate equality of MAD and standard deviation for large, normally distributed samples). Finally, samples with Z_a_ < −5 were removed.

### Differential expression testing

To make differential expression (DE) testing robust against potential outlier measurements (counts) that may remain even after outlier sample removal, we calculated individual observation weights designed to downweigh potential outliers. The weights are constructed separately for each gene. First, Tukey bi-square-like weights (Wilcox, 2012) *λ* are calculated for each (variance-stabilized) observation *x_a_* (index *a* labels samples) as

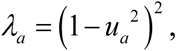

where

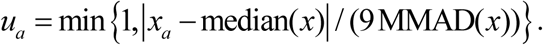

The median is calculated separately for each gene across all samples. MMAD stands for modified MAD, calculated as follows. For each gene, we first set MMAD = MAD. The following conditions are then checked separately for each gene: (1) 10^th^ percentile of the weights λ is at least 0.1 (that is, the proportion of observations with weights <0.1 is less than 10%) (Langfelder & Horvath, 2012) and (2) for each individual genotype, 40^th^ percentile of the weights λ is at least 0.9 (that is, at least 60% of the observation have a weight ≥ 0.9). If both conditions are met, MMAD = MAD. If either condition is not met, MMAD equals the lowest value for which both conditions are met. The rationale is to exclude outliers but ensure that the number of outliers is not too large either overall or in each genotype group. This approach has previously been used in (Lee et al., 2018; Wang et al., 2022).

Preliminary analyses of each separate dataset found that several sets exhibited a relatively large overall variation component unrelated to genotype or sex. Hence, we used the Surrogate Variable Analysis (SVA, R package sva (Leek & Storey, 2007)) to calculate latent factors known as Surrogate Variables (SVs) representing unwanted variation and used the first SV as a covariate in DE testing for these datasets.

DE analysis was carried out using DESeq2 (Love et al., 2014) version 1.44.0 with default arguments except for disabling outlier replacement (since we use weights to downweigh potential outliers) and independent filtering (since we have pre-filtered genes based on expression levels).

### Common analysis of Q140 vs. WT

We collected the Q140 and WT samples from all of our 115 crosses, resulting in a combined set of 902 Q140 and 905 WT samples. To account for batch effects, we tested DE for Q140 vs. WT in each dataset separately using the approach detailed above, including the use of the 1^st^ Surrogate Variable as a covariate where appropriate, and then combined the significance Z statistics for each gene at each age using two approaches. First, we used Stouffer’s meta-analysis method (Stouffer, 1949) to obtain significance statistics that reflect the full number of samples used in the combined analysis. Second, we also combined Z statistics from individual studies using averaging which results in overall significance statistics “scaled” to (approximately) 8 samples per genotype and makes them comparable among the combined studies as well as to DE analyses in individual perturbations. To obtain overall log fold change for each gene, we averaged its log fold changes across all studies.

### Analyses of null perturbations

The large number of WT and Q140 samples generated in this study allowed us to carry out a comprehensive study of DE analysis between two groups that are biologically equivalent (e.g., *n* WT vs. *n* WT or *n* Q140 vs. *n* Q140 samples). Because each KO-het in the study was assayed in a separate batch, we did not sample randomly from the entire set of WT or Q140 samples; rather, we randomly sampled datasets (or KO-hets) and then added a requisite number of samples to both sides of the null DE until we reached the requisite number of samples. We aimed for DE analyses that would have degrees of freedom (DoF) equivalent to *n* = 3, 4, 6, 8, and 10 samples per group without covariates. Because our sampled groups often contained samples from different batches and some batches required the Surrogate Variable covariate, the actual number of samples in each group was somewhat larger than the equivalent *n* so as to result in the same DoF. We carried out 4000 DE analyses for each *n* with groups sampled in this manner (2000 each from WT and Q140 samples). We then calculated transcriptome-wide correlations among the Z statistics for pairs of analyses as well as transcriptome-wide reversal correlations against the common Q140 vs. WT analysis described above. Although the sampled groups are not entirely independent among the analyses, we expect these transcriptome-wide correlations to be a good representation of the null distribution of such correlations. In particular, for the reversal correlations, we find an approximately normal distribution with a standard deviation of about 0.1, and this distribution (and its std. deviation) appears to be essentially independent of the number of samples per group in the range we studied: the standard deviation varies between 0.097-0.098 for 8 and 10 samples per group to 0.112 for 3 samples per group. This suggests that reversal correlation threshold |cor|=0.2 corresponds approximately to a p-value p=0.05 for a null perturbation; we use this value as a suggestive threshold to flag KO-het perturbation for further follow-up. One could also use a more stringent threshold corrected for multiple testing of the 115 perturbations in this study; e.g., a Bonferroni-corrected threshold of p=0.1 corresponds to a reversal correlation of ≈ 0.33.

### Data preprocessing for module preservation analysis

For the analysis of Allelic Series module preservation in the current data, we retained only WT and Q140 samples that passed QC in analyses of individual KO-het datasets. Using these samples as a single unified set, we used the variance stabilizing transformation and then adjusted the data for batch (each individual KO-het is its own batch) as well as the same Surrogate Variables that were used as covariates in the DE analysis of individual KO-het datasets. The adjustment was carried out using empirical Bayes-moderated regression implemented in function empiricalBayesLM in the WGCNA R package.

### Measures of overall transcriptome-wide reversal and exacerbation

We use several statistics to quantify the extent of transcriptomic reversal for each perturbation. These statistics can be broadly divided into two groups, one based on significance and one on fold change. For the former, for each perturbation (KO) we report the numbers of significantly reversed or exacerbated genes at a defined significance threshold (e.g., FDR<0.1), defined as genes that are significantly DE in a baseline Q140 vs. WT test as well as in Q140/KO vs. Q140. Of these, the reversed genes’ fold change has opposite signs in Q140 vs. WT and Q140/KO vs. Q140 (i.e., compared to Q140, expression in Q140/KO moved towards WT levels), while for exacerbated genes the fold changes have the same sign. Although numbers of significantly reversed or exacerbated genes are intuitive, they require selecting a significance threshold and at any given significance threshold they are strongly influenced by the numbers of samples and other technical factors. Another useful statistic is a transcriptome-wide correlation of Z statistics for Q140 vs. WT and Q140 vs. Q140/KO which we call the reversal correlation (of Z statistics). When this correlation is positive (negative), the KO perturbation causes expression changes in Q140 background that moves overall expression levels closer to (further away from) WT levels. Although sample numbers affect the scale of Z statistics, the correlation is scale-independent and hence only sensitive to the number of samples in the sense that higher numbers of samples generally increase signal to noise ratio. One could also define an analogous reversal correlation for log fold changes. Using correlation as a measure of transcriptome-wide reversal is in many ways attractive but it by definition cannot measure the amount of expression reversal or exacerbation, e.g., the fraction of fold change in Q140 vs. WT that is reversed by Q140/KO. Hence, we also defined a measure of overall log fold change reversal. Specifically, we define an “overall reversal fraction” as the coefficient of a weighted linear model regressing log fold changes in Q140 vs. Q140/KO on log fold changes in baseline Q140 vs. WT. The weights are constructed so as to give more weight to genes with stronger evidence of DE in baseline Q140 vs. WT. We chose the form

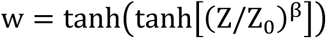

with *Z*_0_=3 and *β=*6. These settings result in weights that are low for *Z* ≤ 3, quickly rise for values around 3 and then flatten out, approaching the asymptotic value of tanh(1) ≈ 0.76 for *Z* much greater than 3. In our applications we find that overall rescue fractions are relatively insensitive to the choices of *Z*_0_ between 1 and 5 and *β* around 6. We further correct the regression coefficient for possible systematic differences between fold changes in Q140 vs. WT in the baseline dataset and in the perturbation study that contains its own Q140 vs. WT test. Using the same weights, we calculate weighted variances of log fold changes for Q140 vs. WT in the baseline and in the perturbation dataset and use their ratio to correct the overall rescue fraction. The rationale for this step is to account for potential dependence of estimated log fold changes on technical variables such as number of samples, biological and technical noise etc.

### Measures of individual gene reversal and exacerbation

For each individual gene, we define two statistics of reversal/exacerbation. First, the reversal score is constructed to identify genes with statistically significant DE for both Q140 vs. WT in baseline data and in Q140/KO vs. Q140. Specifically, it is defined as the minimum of |*Z*_Q140-WT_| and |*Z*_Q140/KO-Q140_|, multiplied by a sign that makes the reversal score positive when the two Z statistics have the opposite signs and negative otherwise; here *Z*_Q140-WT_ and *Z*_Q140/KO-Q140_ are the Z statistics for these two Q140 vs. WT and Q140/KO vs. Q140. Thus, a reversal score of say 4 or more identifies genes that are DE at |*Z*| > 4 in both the baseline and the perturbation test and the DE is such that the perturbation reverses the effect of Q140 back towards WT levels. Since the reversal score is sensitive only to significance but not to the amount of reversal or exacerbation, we also define the reversal fraction as the fraction of the expression change between Q140 and WT that was reversed by the KO perturbation in Q140 background. Specifically, denoting the mean expression levels of a particular gene in WT, Q140 and Q140/KO genotypes by *E*_WT_, *E*_Q140_ and *E*_Q140/KO_, the reversal fraction is defined as *RF* = (*E*_Q140-_ *E*_Q140/KO_)/(*E*_Q140_- *E*_WT_). This can also be expressed in terms of log_2_ fold changes (*lFC*) as

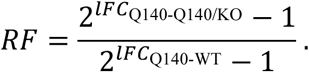

Positive reversal fraction between 0 and 1 means partial correction; reversal fraction is 1 for full correction and above 1 for over-correction. Negative values indicate exacerbation. Together, the reversal score and reversal fraction provide information both about statistical significance and the amount of expression correction. We note that the (individual gene) reversal fractions and (transcriptome-wide) overall reversal fraction, although clearly related, do not measure the reversal on the same scale: The individual gene reversal fraction is based on differences of mean expression values on the natural scale while overall reversal fraction is based on log fold changes, i.e., differences of log-transformed expression values.

Because all of the above reversal measures involve expression in Q140 genotype in two DE comparisons (Q140 vs. WT and Q140 vs. Q140/KO), using the same Q140 data in both comparisons results in a bias towards reversal in both transcriptome-wide and individual reversal statistics. To obtain unbiased reversal statistics, the two DE comparisons need to be carried out on data from independent samples. In this work we use the combined meta-average Q140 vs. WT results as a baseline for reversal analysis. Although the combined Q140 vs. WT results are not, strictly speaking, independent of the samples in each of the individual datasets, we consider the influence of each individual dataset within the meta-analysis or meta-average results small enough to obtain essentially unbiased reversal statistics.

### Measures of perturbation effect on WGCNA modules

Given a WGCNA module, we define the effect of perturbation on the module or the module reversal score as a weighted average of the corresponding gene statistics of the genes in the module. The weight of each gene is chosen to be proportional to the meta-analysis Z statistic of module membership in Allelic Series. Specifically, the Allelic Series modules were defined as consensus modules across three sets; in each set, *Z.kME* for each gene is defined as the Z statistic corresponding to the correlation of the gene expression profile with the module eigengene. Given the three *Z.kME* statistics (one in each set), the Stouffer meta-analysis Z statistic is formed as a sum of the three *Z.kME* statistics divided by 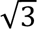. This meta-analysis Z statistic is used as the weight of each gene when forming the module-wide weighted average; the effect is to give higher weight to the hub genes in a module.

This procedure is used both for DE Z statistics, resulting in measures of KO-het effect on modules, as well as for the reversal score which results in module reversal scores. Using our sampling analysis of null perturbations, we are able to evaluate statistical significance of each module statistic. To avoid introducing a (small) bias because the analysis of null perturbations uses a slightly different (smaller) set of genes than each individual KO-het analysis, we restricted the module scores to the genes present in both the null perturbation analysis and the KO-het analysis for which the statistics are calculated.

### Manhattan plots for genes in WGCNA modules

We find it informative to display individual gene DE Z statistics and reversal scores for genes in selected modules in a “Manhattan” plot inspired by plots routinely used to illustrate results of GWAS. Specifically, the plots show DE Z statistics or reversal scores for genes in selected modules which in our application are the 13 strongly CAG-dependent modules in Allelic Series striatum (Langfelder et al., 2016). Genes in each module are ordered from left to right by decreasing module membership, specifically the meta-analysis *Z.kME* statistic. A local weighted average of the statistic is shown as an orange line that indicates the overall trend of the statistic. The weight of gene *j* in the local average at position *i* are proportional to 𝑒𝑥𝑝((𝑖 − 𝑗)^2^/(2 ∗ 50^2^)), i.e., are Gaussian with standard deviation of 50. A barplot below the scatterplot shows a weighted average of the displayed statistic with weights that are proportional to the module membership meta-analysis *Z.kME* statistics. Genes that pass a certain threshold, e.g., FDR<0.1 for DE Z statistics, are shown in red or blue depending on the sign of their statistics.

### Enrichment analysis

Enrichment analyses were performed using our published methods (Wang et al., 2022). Briefly, enrichment calculations were carried out on sets of genes DE at FDR<0.1 or p<0.01 using our in-house R package anRichment (https://github.com/plangfelder/anRichment) that implements standard Fisher exact test and a multiple-testing correction across all query and refence gene sets. To quantify the reversal/exacerbation effect of KO perturbations on Allelic Series WGCNA modules, we tested enrichment of genes DE in Q140/KO vs. Q140.

### Clustering of perturbations based on enrichment of DE genes

We clustered perturbations and enrichment terms by the enrichment ratio, defined as the ratio of observed and expected overlap of the corresponding set of DE genes with a literature set. We manually selected most informative and relevant gene sets from the multitude of often very similar literature sets.

### Weighted transcriptome-wide correlations as a measure of similarity among effects of KO- hets

Although one can correlate transcriptome-wide Z statistics of different KO-het perturbations using standard Pearson correlation, we found that weighted correlations tend to be more informative when genes strongly DE in Q140 vs. WT carry a higher weight. Specifically, the weights are proportional to the DE Z statistics in our common analysis of Q140 vs. WT across all datasets.

### Statistical analysis

Unless otherwise noted, data are presented as mean ± standard error of the mean (SEM) and statistical significance of differences was assessed using ANOVA followed by Tukey HSD test. We have not tested the residual errors for normality. Numbers of samples are indicated in text and in figure legends. Differential analysis details, including down-weighting of suspected outliers, are described in Method Details section. Enrichment was evaluated using the hypergeometric test (also known as Fisher’s exact test) with the set of analyzed genes (after filtering out low-expressed ones) used as the background. Data were analyzed using R versions 4.1.1, 4.2.1 and 4.4.1, and GraphPad Prism 7.

## ACKNOWLEDGEMENT

The research was fully supported by the CHDI Foundation, Inc. X.W.Y is also supported by the Terry Semel Chair in Alzheimer’s Research and Treatment at UCLA, and by generous donations from HD families and their supporters. We thank Giovanni Coppola for his inputs in developing the Manhattan plots for genes in the AS CAG-dependent WGCNA modules. We also thank UCLA Neuroscience Genomics Core (UNGC), UCLA Technology Center for Genomics & Bioinformatics (TCGB), the Jackson Laboratories, Mutant Mouse Resource & Research Centers (MMRRC), the European Mouse Mutant Archive (EMMA), the NCI Mouse Repository, Experimental Animal Division (RIKEN BRC) and Rodent Genetics Core at Cedar-Sinai for their research support.

## AUTHORS CONTRIBUTIONS

X.W.Y. conceptualized, designed, and supervised the study, and X.W.Y., P.L. and N.W. wrote the manuscript. X.W.Y., N.W., P.L., S.H., G.C., J.R., J.A. and T.V. participated in data interpretation. N.W. led a group of researchers, L.R., X.G., M.S., M.P., R.V. and J.R. to perform the mouse model generation/import, colony management, tissue dissection and RNA-seq studies. F.G. and P.L. performed RNA-seq data processing. P.L. performed all the bioinformatic analysis and generated relevant plots. N.W. performed pathological studies. Y.O., S.L. and A.Y. performed Scn4b and Kcnh4 perturbation study in patient-derived MSNs.

## DECLARATION OF INTERESTS

X.W.Y. is a current or former Scientific Advisory Board member for Lyterian and Ophidion, and has served as a consultant for Ionis, Biogen, Novartis, Roche, LifeEdit, PTC, Ascidian, Sangamo, Forbion, and Harness. P.L. serves as occasional bioinformatics consultant for The Bioinformatics CRO, Inc; Vynance Technologies, LLC; and FOXO Technologies, Inc. M.S. is a full-time employee at, and holds shares in, Alexion Pharmaceuticals.

---

**Supplemental Figure S1.**
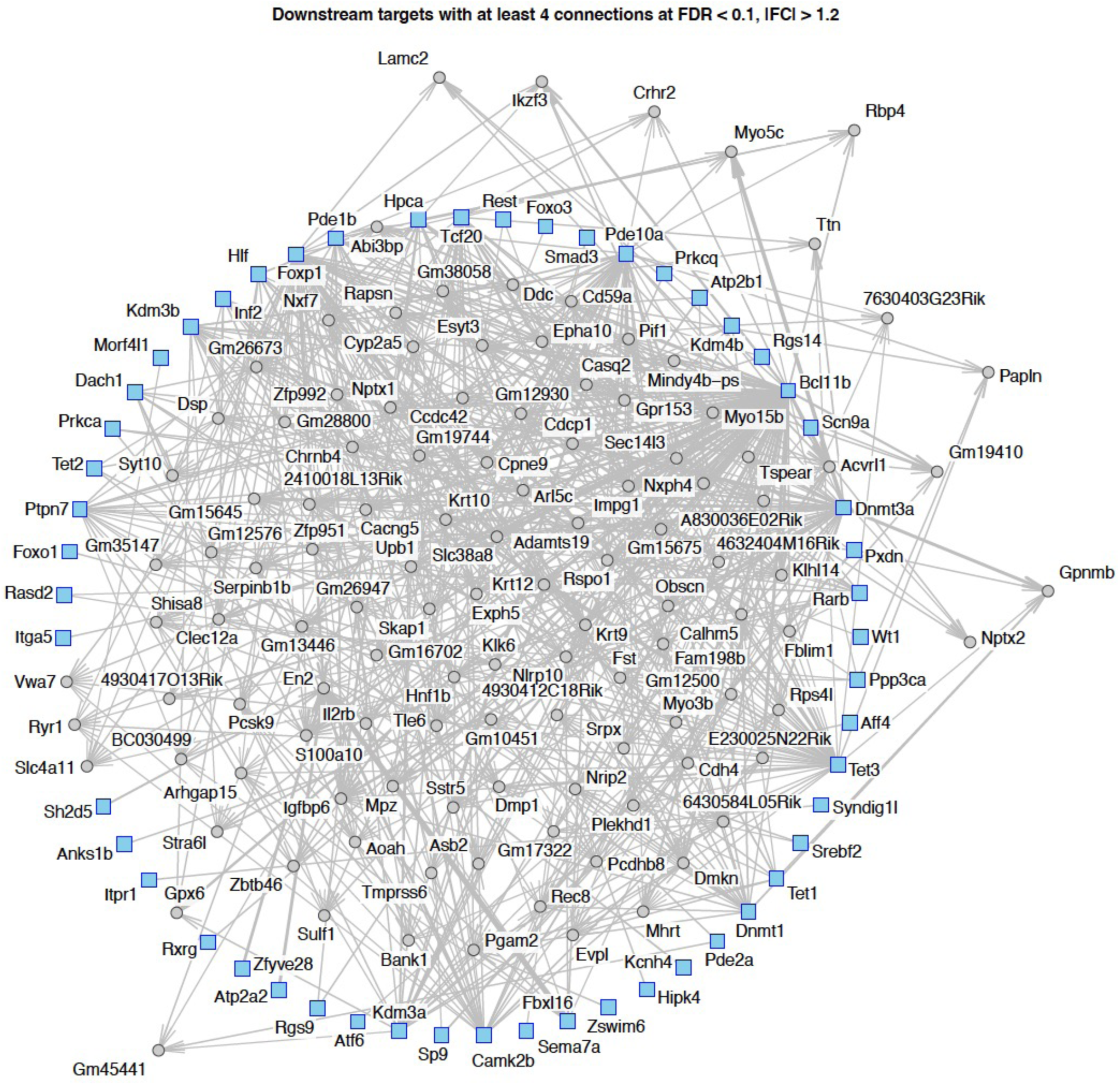
Network of KO-het perturbations and downstream genes. Network plot of our 115 perturbations and genes affected by the perturbations in WT background. Only those downstream genes are shown that are affected by at least 4 perturbations at FDR<0.1 and fold change |FC|>1.2

**Supplemental Figure S2.**
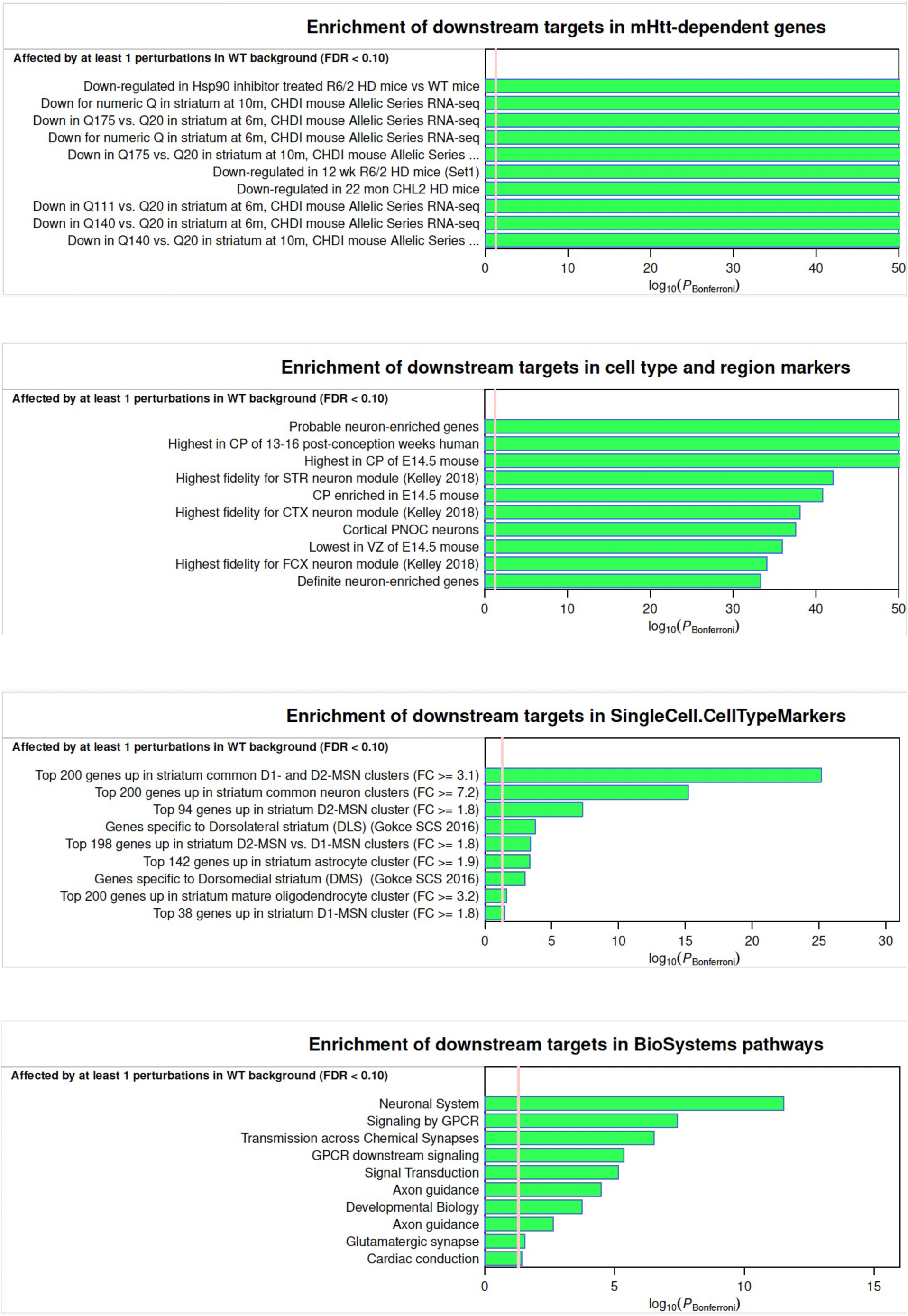
Enrichment analyses of dysregulated genes by at least one perturbation. Bonferroni-corrected enrichment significance of the set of genes affected at FDR<0.1 by at least one perturbation in WT background. Each panel shows the top 10 terms in one collection of reference gene sets.

**Supplemental Figure S3.**
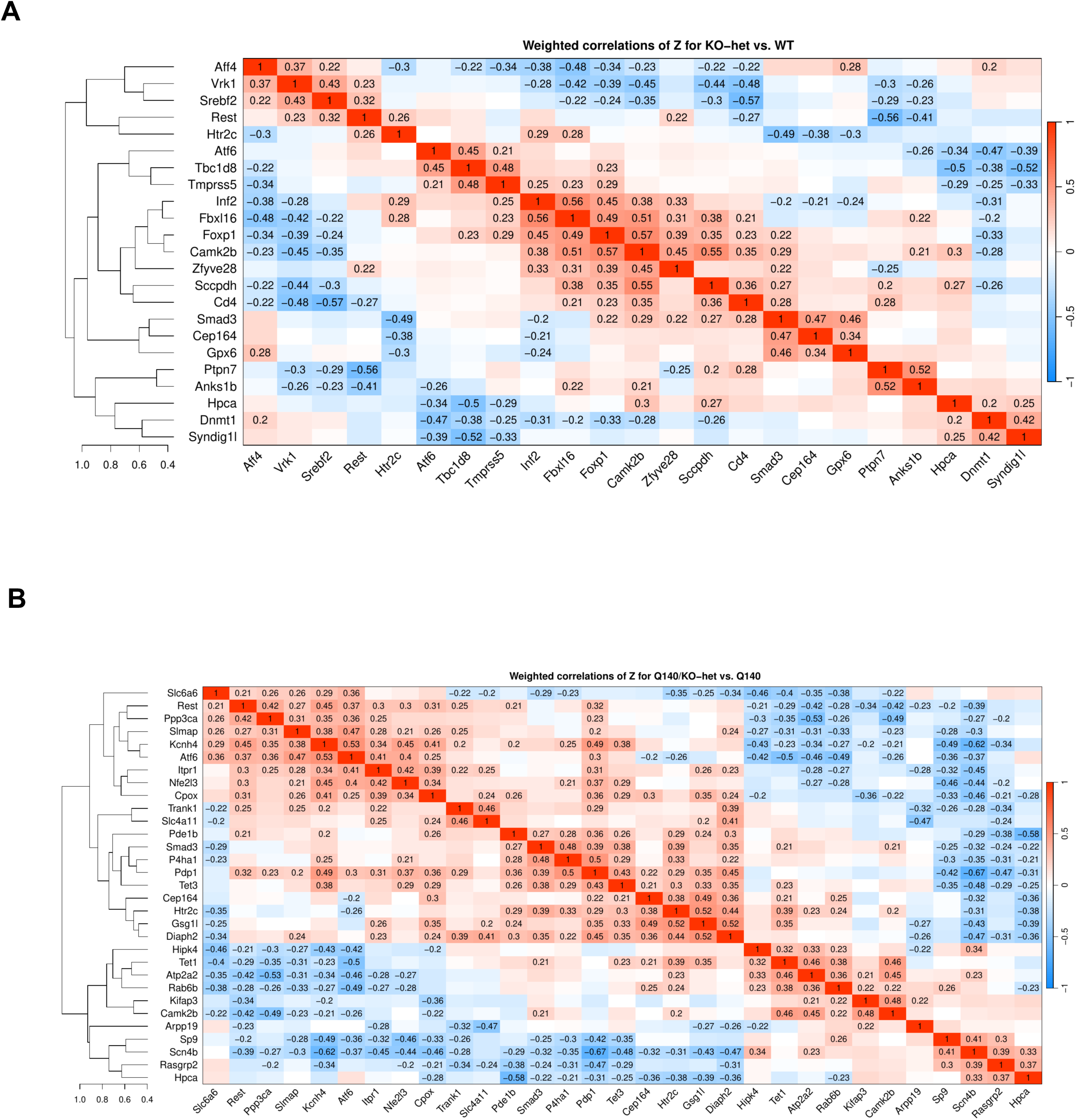
Correlations among perturbations in WT or Q140 background. (A) Heatmap representation of weighted correlations of Z for KO-het vs. WT for selected perturbations, namely those that have at least one correlation with another perturbation whose absolute value is at least 0.45. Correlations whose absolute value is above 0.2 are shown explicitly. Average-linkage hierarchical clustering tree is shown on the left. (B) Analogous heatmap of weighted correlations of Z for Q140/KO-het vs. Q140.

**Supplemental Figure S4.**
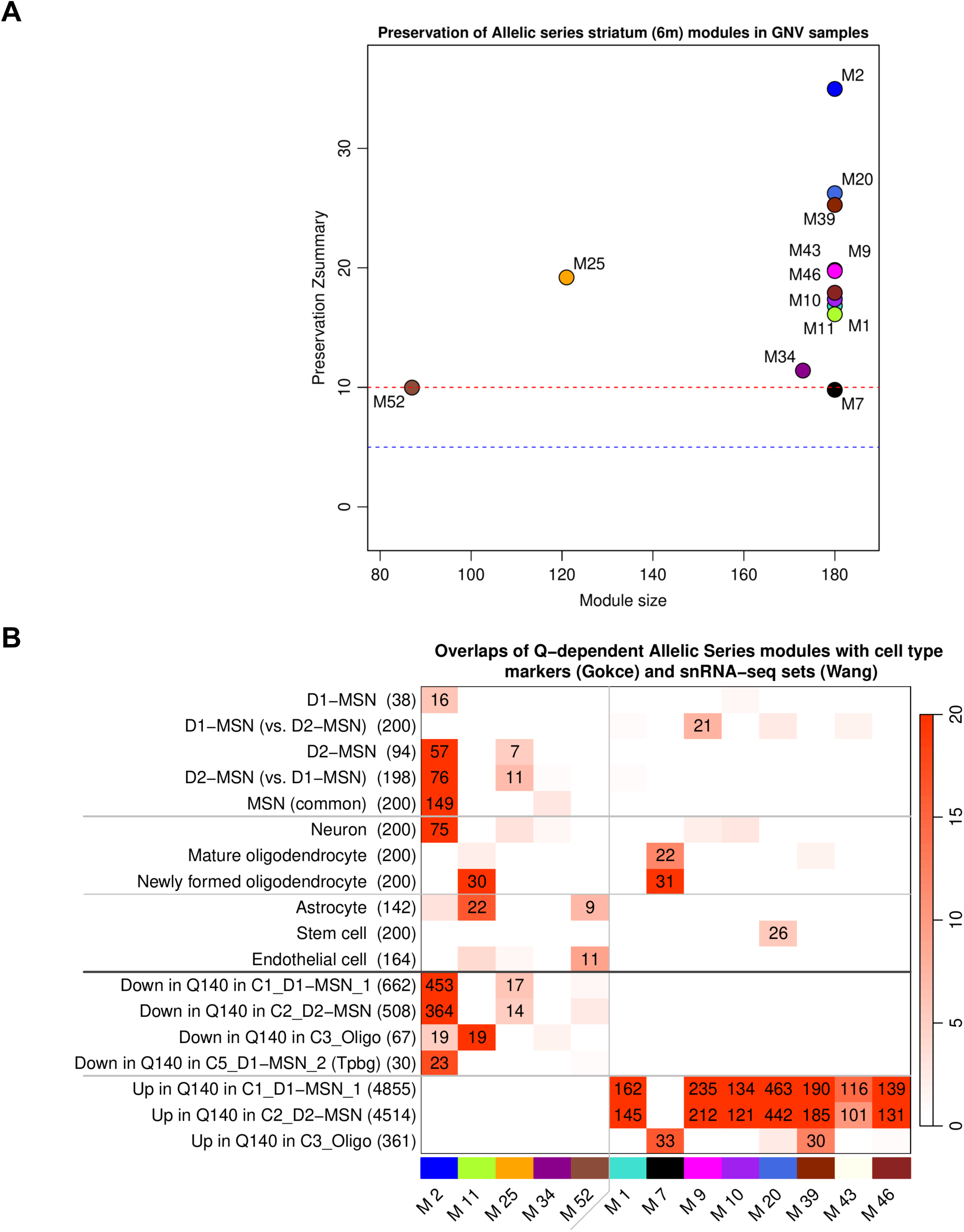
Preservations and cell type enrichment of Allelic series CAG-dependent modules. (A) Preservation Zsummary statistics for the 13 strongly CAG-dependent Allelic Series modules in Q140 and WT data from this study. Module size displayed on the x-axis reflects the lesser of 100 and the number of genes in each module which is the effective size of the module in preservation calculations. Blue and red dashed lines represent the thresholds for moderate and strong overall evidence of preservation, respectively. (B) Enrichment of CAG-dependent Allelic Series striatum modules in cell type marker sets distilled from single cell RNA-seq of mouse striatum and in genes DE for Q140 vs. WT in individual clusters in snRNA-seq analysis of Q140 and WT striatum.

**Supplemental Figure S5.**
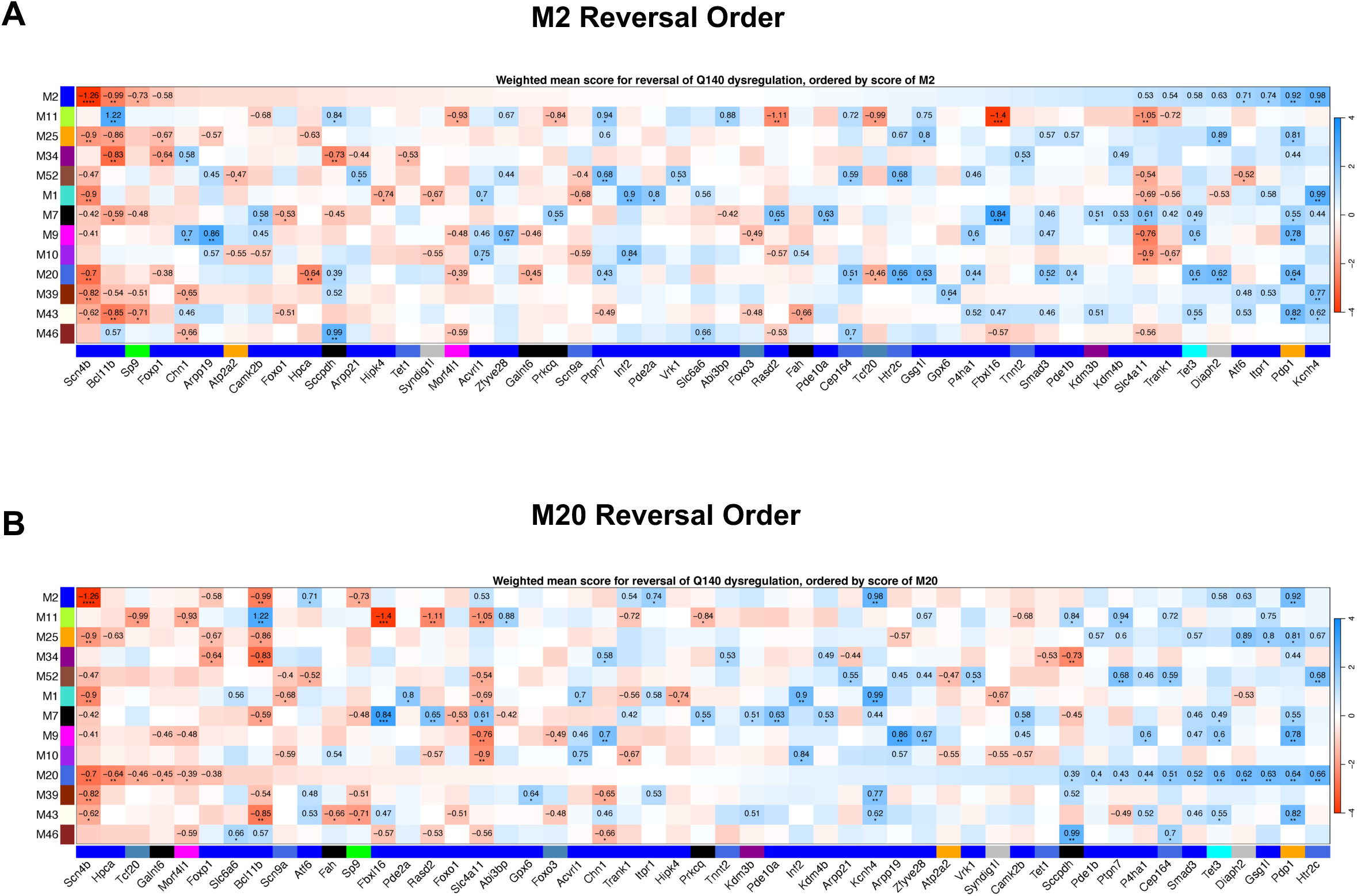
Ranking of GNV perturbations on their reversal effects of M2 or M20 modules. (A) Heatmap representation of weighted mean module reversal scores. Scores with corresponding (nominal) p-value below 0.1 are shown as numbers. Stars indicate nominal significance: *, p<0.05; **: p<0.01; ***: p<0.001. Only those perturbations are shown that affect at least one module at a nominal p<0.05. Perturbations are ordered by increasing reversal score of M2. (B) The same data as in (A) but perturbations are ordered by increasing reversal score of M20.

**Supplemental Figure S6.**
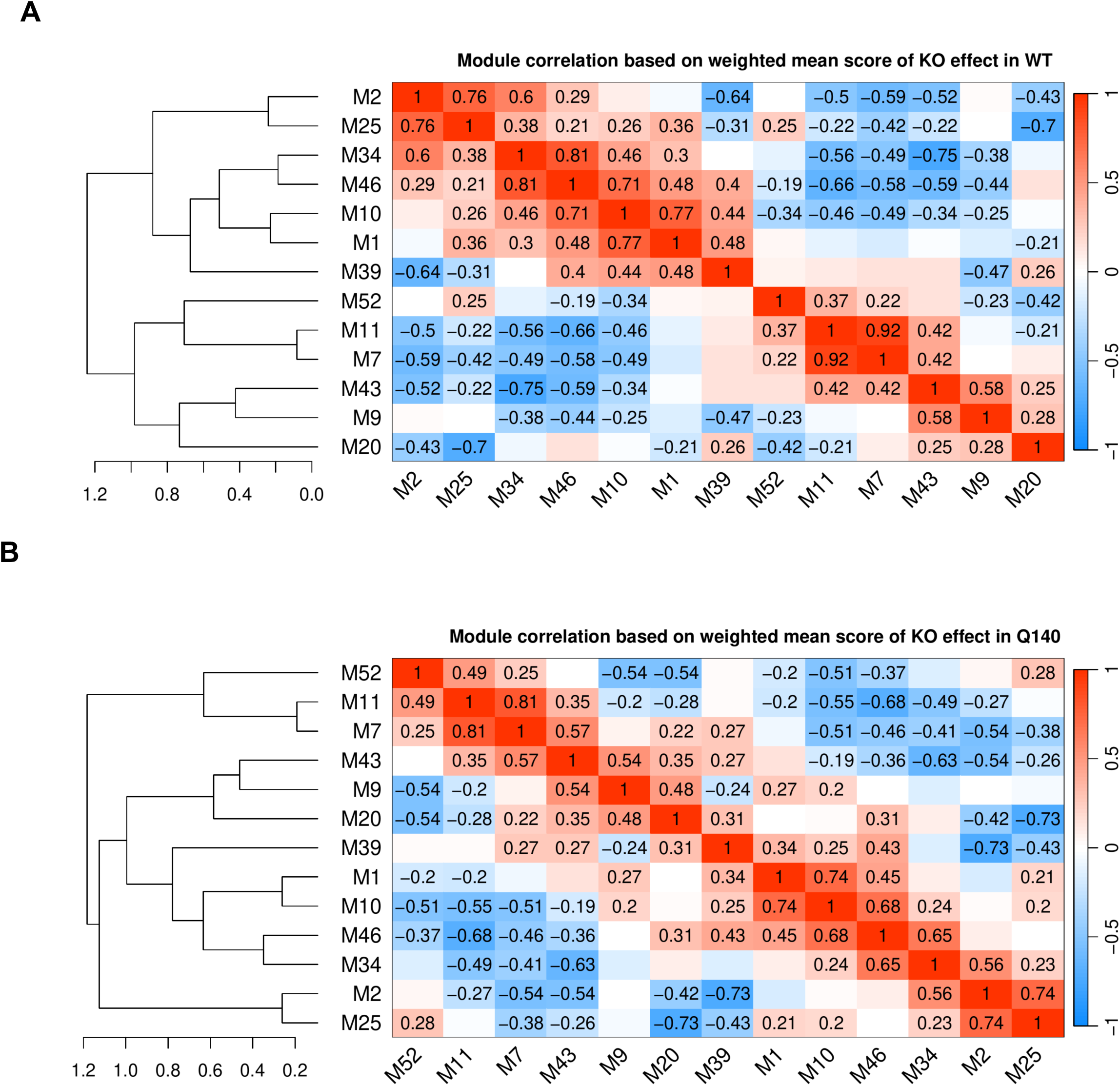
Correlations of the CAG-dependent modules based on KO-het effects. (A) Correlation heatmap of module effect scores of KO-het vs. WT in the 13 strongly CAG-dependent Allelic Series modules. Entries that correspond to nominal p-values p<0.05 are shown explicitly. Average-linkage hierarchical clustering tree is shown on the left side of the plot. (B) Analogous heatmap of module effect scores of Q140/KO-het vs. Q140.

**Supplemental Figure S7.**
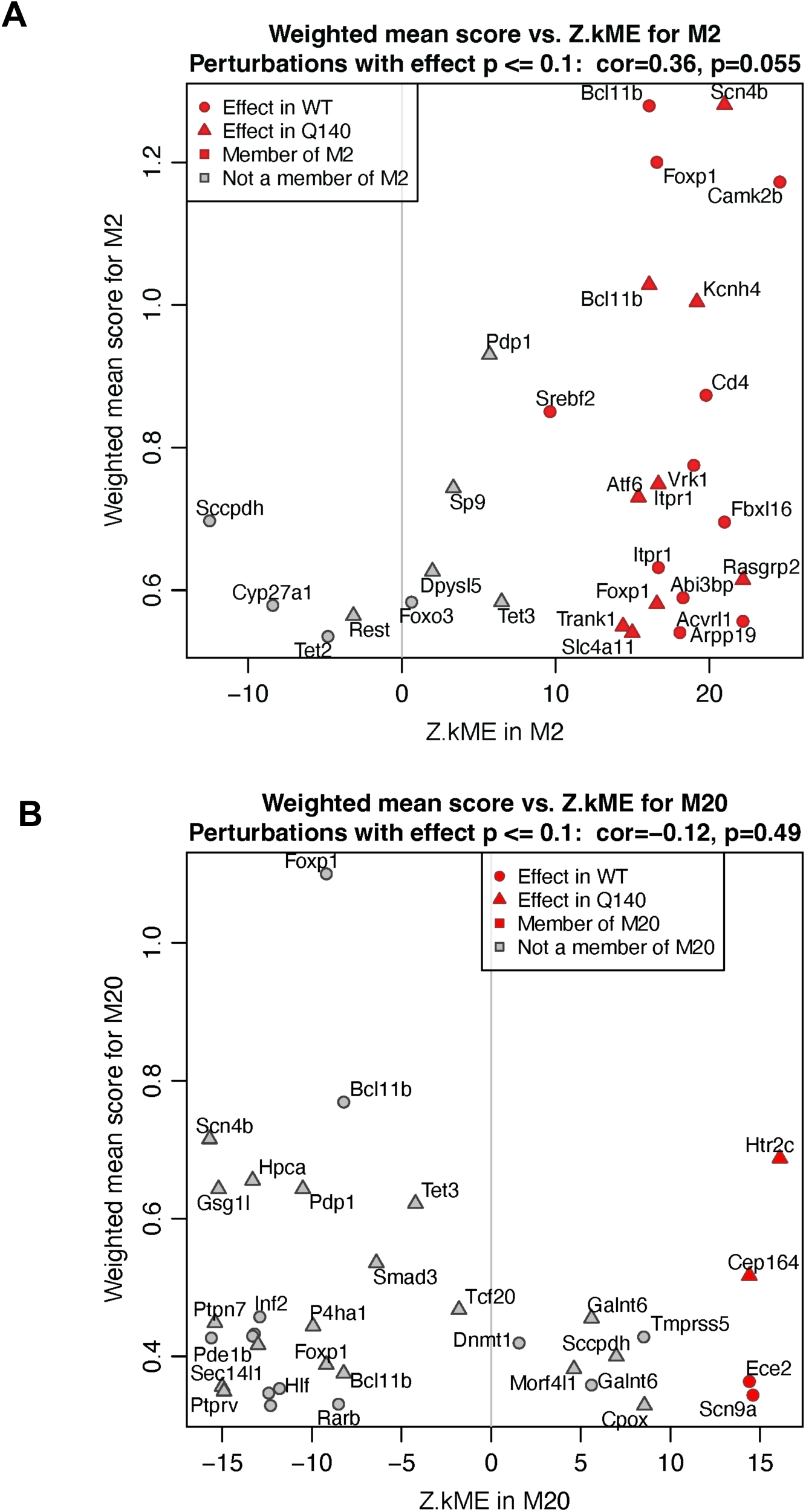
Relationships of GNV gene module memberships and their perturbation effects in M2 and M20. (A) Scatterplot of Allelic Series module M2 weighted mean effect scores of KO-het vs. WT (circles) and Q140/KO-het vs. Q140 (triangles) against the module membership measure Z.kME. Red color denotes members of module M2. Only those perturbations are present in the plot that show an effect with nominal p-values less than 0.1. (B) Analogous scatterplot for effect scores of module M20.

**Supplemental Figure S8.**
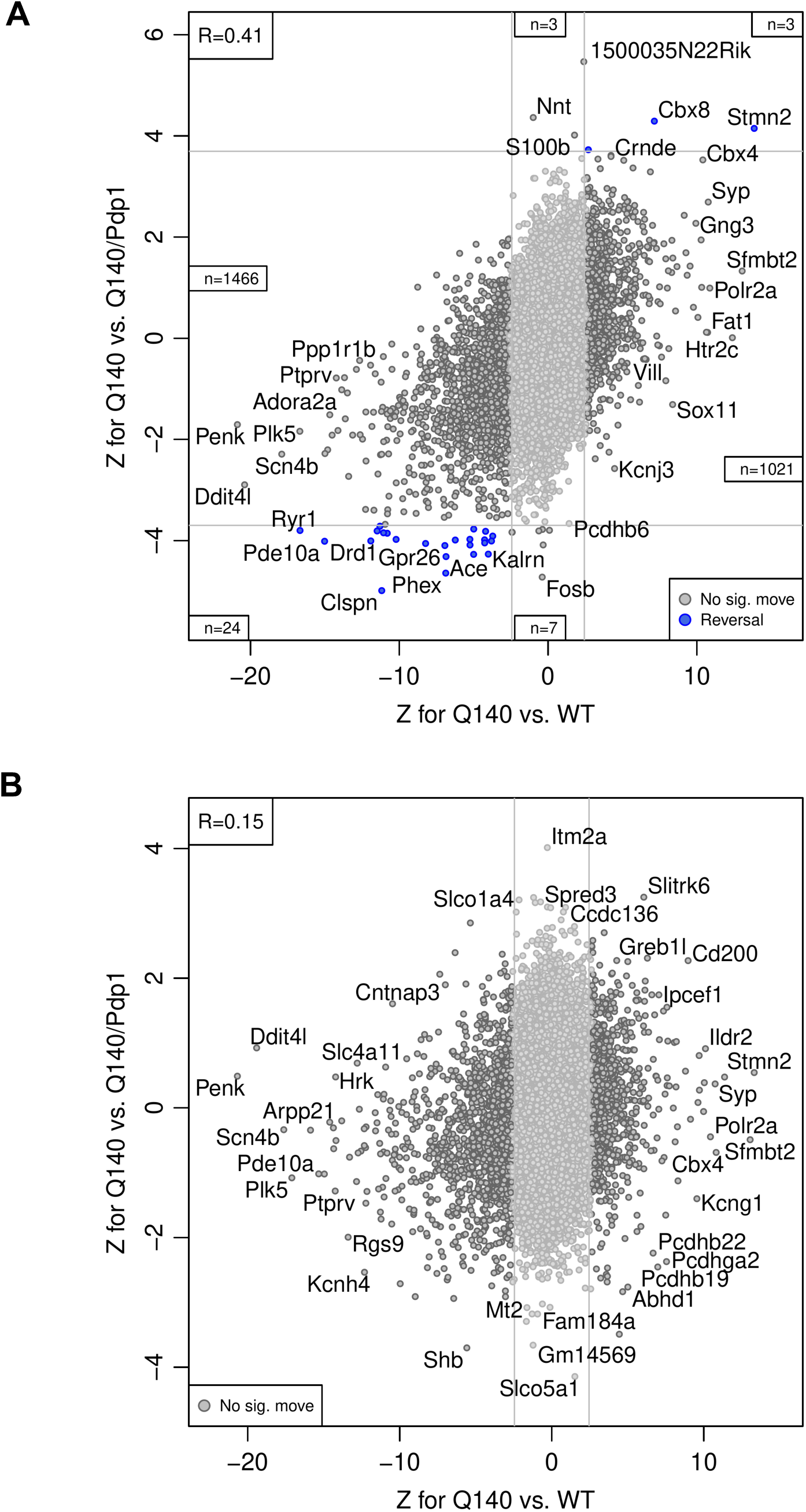
Genome-wide reversal scatterplots of Pdp1/Q140 by sex. Reversal scatterplot for Pdp1 in female (A) and male (B) samples. X-axis shows Z for Q140 vs. WT in our common analysis of all Q140 and WT samples, while the y-axis shows Z for Q140 vs. Q140/Pdp1. Grey lines indicate approximate locations of FDR=0.1 threshold. Blue and red color indicate genes reversed and exacerbated, respectively, at FDR<0.1.

